# Synthetic Cytosolic Splicing Enables Programmable mRNA-Encoded Receptors

**DOI:** 10.1101/2025.04.27.650769

**Authors:** Matteo Lampis, Yaakov Benenson

## Abstract

A typical cell therapy product comprises engineered cells that detect disease-related molecular cues in their surrounding or on a surface of a target cell, and activate a response that alleviates disease symptoms or eradicates diseased cells. mRNA as a therapeutic substrate has become prevalent in the last decade across multiple therapeutic areas, and it has also been evaluated as a building block of cell therapies. However, compared to DNA-based building blocks, it is much more challenging to use mRNA in a programmable manner to engineer complex multi-input/multi-output processes that can fully support the next generation of cell and gene therapies. Addressing this challenge requires the exploration of novel post-transcriptional control mechanisms that bridge mRNA regulation with extracellular surroundings.

Here, we engineer a family of **s**ynthetic **m**RNA **s**plicing (SMS) receptors by redesigning the Inositol-requiring enzyme 1 (IRE1) to regulate protein synthesis from a precursor mRNA. We design SMS-based receptors that sense diverse intracellular and extracellular inputs, highlighting the versatility and modularity of this platform. We apply this approach to design a ‘cytokine-converter’ receptor that detects inflammatory cytokines and produces an anti-inflammatory output in response. That receptor is successfully validated in cell lines and primary T cells upon mRNA delivery. These cells generate anti-inflammatory IL-10 upon stimulation by physiological levels of either TNF-α or IL-1β secreted by macrophage-like cells, highlighting their potential as a cell therapy for inflammatory diseases. With its modular and programmable architecture, the SMS platform is poised to become an important enabling tool for sophisticated programmable mRNA therapeutics.

## Introduction

With the success of COVID-19 vaccines, messenger RNA (mRNA) has emerged as an efficacious and safe therapeutic modality with billions of vaccine doses administered globally^1^. Expanding mRNA-based technologies beyond vaccination could drive breakthroughs in treating other diseases, enabling a transient delivery of genetic instructions with reduced risk of insertional mutagenesis^2, 3^ and epigenetic silencing^4^, while also lowering manufacturing cost^5^. However, while mRNA holds great clinical potential, its ability to encode complex regulatory mechanisms remains limited^6^. In contrast, DNA-based systems have already been extensively engineered^7^, resulting in sophisticated synthetic gene circuits capable of conditional and dynamic multi-input information processing. Initially, protein expression from DNA constructs utilized constitutive promoters^8^, producing proteins in a poorly controlled fashion. The development of regulated transcriptional systems, such as small molecule-inducible promoters^9^, introduced conditional expression in response to environmental stimuli. These advances ultimately led to the engineering of synthetic receptors^10^ that couple extracellular ligand recognition to transcriptional activation, enabling programmable control over cellular functions. For example, SynNotch^11^, MESA^12^, and Tango systems^13, 14^ involve fusing a transcription factor to the intracellular part of the receptor and releasing it through single-turnover proteolytic cleavage upon a stimulus. Other strategies, such as GEMS^15^ and the artificial two-component system^16^, employ an orthogonal transcription factor activated through multiple-turnover receptor-mediated phosphorylation, allowing for sustained gene expression upon receptor engagement. These receptor-based strategies have enabled the design of highly customizable cell therapy prototypes, for example with the production of a CAR precisely controlled by multiple cell surface tumor associated antigens^17^. While DNA-based synthetic receptors provide a modular framework for engineered cellular responses, mRNA technologies still lack the components required for programmable sensing of extracellular signals, because current RNA-regulation platforms^18, 19^ respond exclusively to intracellular ligands. For instance, one approach to achieving conditional protein expression from an mRNA template is the incorporation of microRNA target site into engineered transcripts, enabling cell-type-specific repression of protein translation based on endogenous microRNA levels^20^. Additional developments in RNA-based regulation include ligand-dependent control using toehold switches^21^ and riboswitches^22^, which enable intracellular ligands and RNA molecules to trigger protein synthesis. Ribozymes^23^ and aptamers^24^ can also regulate RNA stability through cleavage when combined into synthetic regulatory architectures^25^. Expanding upon RNA-acting RNAs mechanisms, which rely on intrinsic sequence-encoded interactions, protein-based RNA regulators introduced an additional and modular layer of post-transcriptional control. For instance, orthogonal proteases have been designed to cleave RNA-binding proteins or activate RNA-modifying enzymes in response to intracellular stimuli^26^, offering programmable post-transcriptional regulation. Finally, adenosine deaminases acting on RNA systems^27, 28^, have been engineered to function as programmable RNA editors, sensing intracellular mRNA inputs to mediate targeted adenosine-to-inosine modifications. However, despite significant progress in modulating RNA activity with intracellular inputs, the field still lacks the tools capable of transducing extracellular signals using exclusively the mRNA and mRNA-encoded proteins, but without the use of DNA components.

To address this gap, here we present an engineered modular post-transcriptional synthetic receptor that directly couples the presence of an extracellular or an intracellular ligand to protein translation from an mRNA template based on the transduction mechanism of the inositol-requiring enzyme-1 (IRE1)^29, 30^. IRE1 is an evolutionary conserved single-pass transmembrane protein that resides in the endoplasmic reticulum (ER) of all eukaryotic cells^31^ with the capacity of performing inducible splicing of cytosolic mRNA^32^. Upon homodimerization induced by ER stress, IRE1 activates its cytosolic ribonuclease (RNase) domain^33^, which cleaves a consensus sequence within the mRNA encoding the x-box binding protein 1 (XBP1) in mammals^34^. The net effect of this cleavage is the excision of the XBP1 internal cytosolic intron^35^ which induces a translational frameshift that produces the longer and active XBP1 form, a transcription factor that regulates the unfolded protein response genes to counteract the ER stress^36^. Previous works suggest that this cytosolic mRNA splicing pathway could be further engineered^37^, as purified IRE1’s kinase-RNase domains process RNA independently of luminal and transmembrane regions^38^, while the XBP1 splicing hairpin remains functional when transplanted into exogenous RNAs^39, 40^.

Inspired by the mechanism of action of IRE1, and borrowing certain component of this pathway, we engineered custom **s**ynthetic **m**RNA **s**plicing (**SMS**) pathways by triggering the IRE1 kinase-RNase domain dimerization via their fusion to extracellular recognition domains, including nanobodies and natural ectodomains, enabling the detection of both soluble and cell-surface antigens. To prevent the activation of receptor output with the endogenous unfolded protein response pathway via human IRE1, we designed our receptors using orthologous mRNA hairpins that are processed by non-human, engineered IRE1 cytosolic splicing domains. Importantly, we also demonstrate that our synthetic receptors can be efficiently activated when delivered solely as mRNA to different human cell lines and primary T cells.

## Results

### Reprogramming of mRNA Cytosolic Splicing Using Human-Derived IRE1 Intracellular Domains

In the endogenous unfolded protein response pathway, accumulation of misfolded proteins into the ER during stress triggers the homodimerization of the IRE1 luminal domain^41^, forcing the two cytosolic kinase domains to juxtapose in a transphosphorylation-competent orientation (Fig. 1a). The phosphate addition in the activation loop of the kinase region induces a conformational change that allows the two RNase subunits to form the active catalytic site, which cleaves a species-specific consensus sequence^42^ present in tandem on a bifurcate stem loop hairpin of the XBP1 mRNA. The dual cut performed by the IRE1 RNase allows the subsequent rearrangement of the mRNA secondary structure, which provokes the ejection of the internal 26-base cytosolic intron fragment^35^, followed by the sealing of the remaining scar through the RtcB tRNA ligase^43^.

**Figure 1.**
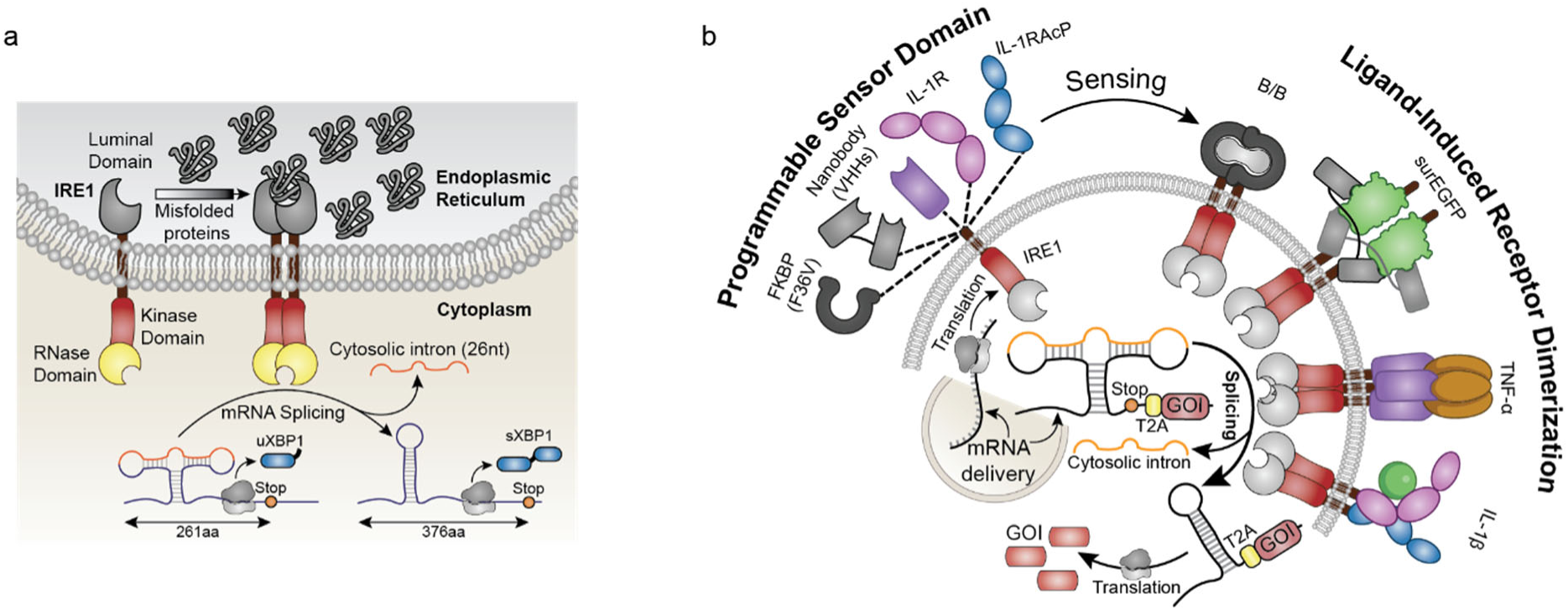
Design of programmable mRNA-only SMS receptors. **a,** Schematic illustration of the endogenous IRE1 pathway. IRE1 is an ER-resident transmembrane receptor with luminal and cytosolic domains. Under stress conditions, misfolded proteins accumulate in the ER, leading to the dimerization of the luminal IRE1 domain and the activation of the cytosolic domain. This enables the RNase domain of IRE1 to splice a 26-nucleotide intron from unspliced XBP1 (uXBP1) mRNA, resulting in the production of spliced XBP1 (sXBP1), a key transcription factor in the unfolded protein response. The unspliced XBP1 (uXBP1) produces a 261-amino acid (aa) product, while the frameshift in the spliced XBP1 (sXBP1) results in a longer 376-aa product. **b,** Engineering of the SMS system for programmable mRNA splicing. The platform is delivered as an mRNA encoding the programmable receptor and the synthetic splicing target. The receptor consists of ligand-induced actuator domains derived from the kinase-RNase domains of IRE1, and a modular extracellular ligand-binding domain (such as FKBP, nanobodies, IL-1R/IL-1RAcP) for detecting their cognate soluble ligands (such as B/B, surface EGFP (surEGFP), IL-1β, TNF-α). Upon ligand binding, the SMS receptor undergoes dimerization, activating the IRE1 RNase domain, which results in the splicing of the cytosolic intron from the synthetic target mRNA. This splicing leads to the translation of the gene of interest (GOI) downstream of a T2A self-cleaving peptide, ensuring the independent translation of the desired protein product from the spliced hairpin.

We hypothesized that this dimerization-induced cytosolic splicing mechanism could be harnessed to develop a platform for post-transcriptional control of any arbitrary protein-coding mRNA (Fig. 1b).

To monitor the IRE1 receptor activity, we adapted a previously developed^44^ reporter of the unfolded protein response, by inserting the XBP1 splicing region (nucleotides 410-633) upstream of the out-of-frame coding sequence of the bright monomeric red fluorescent protein mScarlet (Fig. 2a). The removal of the cytosolic intron by IRE1 leads to a frameshift, restoring the translation of mScarlet, thereby coupling the splicing activity of the receptor to a fluorescent signal. To evaluate the dynamic range of our splicing reporter, we transfected the reporter-encoding DNA plasmid into HEK293 cells treated with tunicamycin, an ER-stress inducer that activates IRE1^45^. This resulted in an 11-fold fluorescence increase compared to controls (Fig. 2b). RT-PCR confirmed intron removal within 2 hours (Fig. 2c, 2d), with residual splicing in untreated cells likely due to transfection-induced stress (Supp. Fig. 1a-c). To rule out leaky translation, we tested the reporter in IRE1 knockout HeLa cells, which showed no fluorescence signal (Fig. 2d, 2e), and therefore we performed further characterization in this knockout cell line to avoid interference from endogenous IRE1.

**Figure 2.**
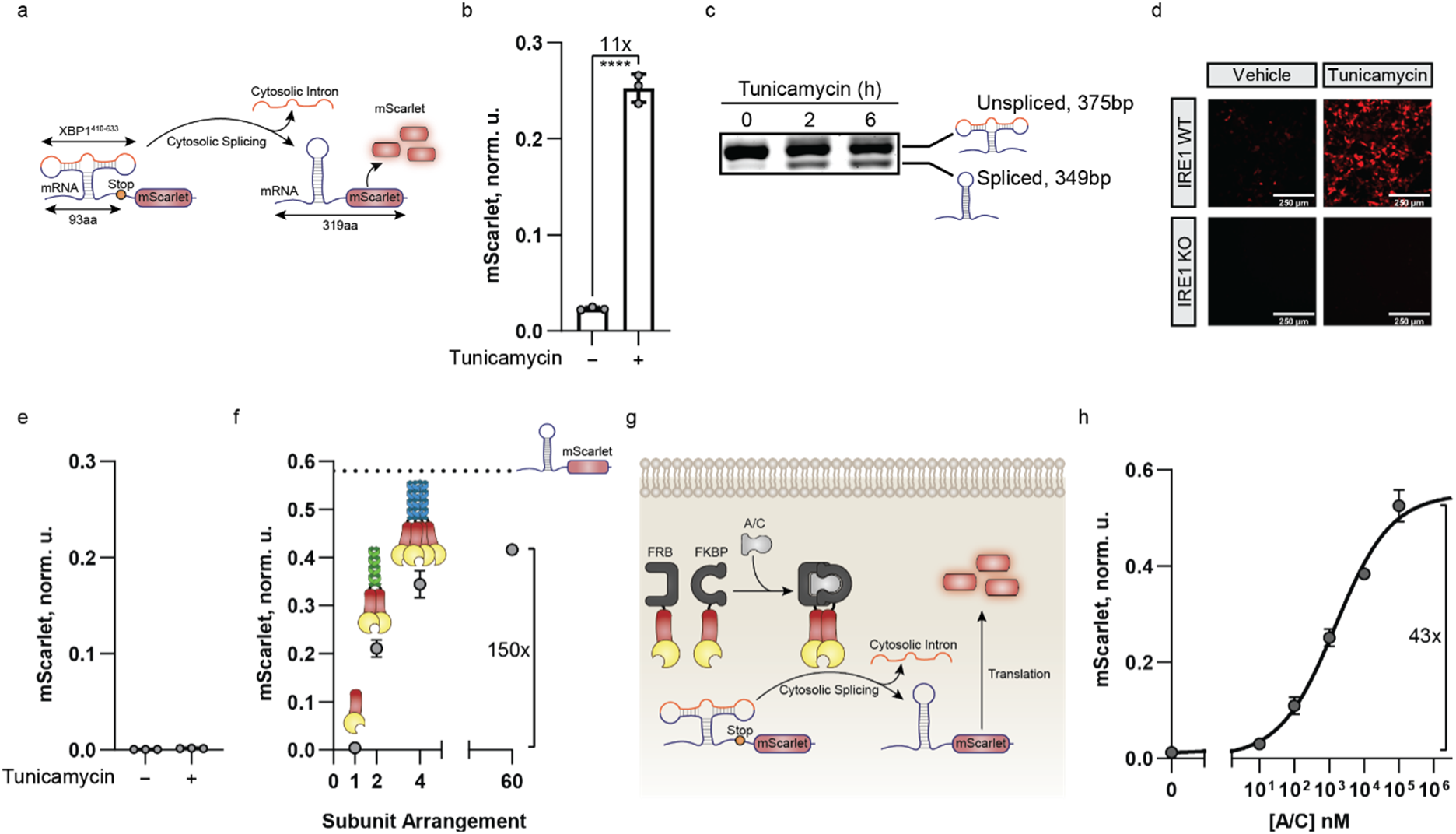
Reprogramming Cytosolic mRNA Splicing Using Human-Derived IRE1 Domains. **a,** Schematic representation of the mRNA splicing reporter. The reporter consists of the XBP1 mRNA splicing region (nucleotides 410-633) inserted upstream of the out-of-frame coding sequence of the fluorescent protein mScarlet. Upon removal of the cytosolic intron by IRE1, the translation of mScarlet is restored, providing a fluorescent readout of the receptor splicing activity. **b,** Splicing reporter output in HEK293 cells treated with the ER-stress inducer tunicamycin. The bar chart shows the mScarlet normalized units (norm. u.) for biological triplicates, n=3. Individual data points are overlaid in gray, with error bars indicating the standard deviation. An unpaired two-tailed t-test was performed, and asterisks indicate p < 0.0001 (****). **c,** Semi-quantitative RT-PCR gel showing the unspliced (top, 375 base pairs) and spliced (bottom, 349 base pairs) forms of the splicing reporter in HEK293 cells treated with tunicamycin for 0 (control), 2, or 6 hours. **d,** Fluorescence microscopy images of HEK293 or HeLa IRE1 knockout (KO) cells transfected with the mRNA splicing reporter and treated with tunicamycin to induce ER stress. Increased red fluorescence indicates splicing activity of the IRE1 receptor. Scale bars represent 250 μm. **e,** Splicing reporter output in HeLa IRE1 KO cells upon addition of tunicamycin. The bar graph illustrates the normalized units of mScarlet (norm. u.) for three biological replicates (n=3). Gray points denote individual data, with error bars indicating the standard deviation. **f,** Splicing reporter induction with the human IRE1 cytosolic fragment fused with constitutive self-assembly domains in HeLa IRE1 KO cells. The fluorescent mScarlet (norm. u.) signal increases with dimerization stoichiometry for biological duplicates (n=2). The dashed line indicates the output of a genetically encoded, constitutively spliced fluorescent reporter driven by the EF1alpha promoter. **g,** Schematic of the inducible heterodimerization system using FKBP and FRB domains fused to IRE1 cytosolic regions. The addition of the small molecule A/C induces heterodimerization, activating the IRE1 splicing domains and inducing the production of mScarlet by the reporter. **h,** Dose-response curve showing the normalized mScarlet fluorescence in response to varying concentrations of the heterodimerizer A/C. Each data point shown is the average of biological triplicates (n=3), with error bars indicating the standard deviation.

As a first step in designing a post-transcriptional sensor, we explored whether the cytosolic domains of IRE1 could be reprogrammed to dimerize and initiate mRNA splicing independently of their native transmembrane and luminal ER-sensing regions. Given previous studies suggesting that higher-order oligomerization of IRE1 may be important for its activation^46^, we sought to determine whether dimerization alone was sufficient to induce splicing of our synthetic reporter. To test this, we fused the human IRE1 cytosolic fragment (IRE1^467–977^) to well-characterized constitutive oligomerization domains with defined subunit arrangements^47–49^, ranging from dimerization to a 60-subunit dodecahedron. Although the splicing reporter showed signal activation that correlated with the oligomerization stoichiometry (Fig. 2f), with the dodecahedron achieving a 150-fold increase compared to the cytosolic IRE1 fragment, dimerization alone was sufficient to induce a 50-fold increase. This result demonstrates that in this setting, dimerization is an effective strategy to induce cytosolic splicing. Notably, the mRNA splicing reaction achieved 75% of the signal generated by a positive control lacking the synthetic cytosolic intron driven by the strong EF1alpha promoter, indicating its efficiency in producing high levels of outputs (Fig. 2f). Additionally, we verified the potential to control the cytosolic splicing reaction by a non-native ligand, a crucial feature for any programmable sensing device. To this end, we fused IRE1^467-977^ to the C-terminus of each subunit of a well-characterized inducible heterodimer^50^ (Fig. 2g), composed of the FK506-binding protein 12 (FKBP) and FKBP12-rapamycin binding domain^T2098L^ (FRB^T2098L^). The dose-response curve (Fig. 2h) with varying concentrations of the small molecule heterodimerizer A/C exhibited the expected Hill function behavior with an EC50 of 15 nM and a dynamic range of 43-fold. These results collectively demonstrated that cytosolic splicing can be triggered either through forced dimerization or finely tuned in a dose-dependent manner via a small molecule ligand.

### Development of SMS: an Orthologous IRE1-Hairpin Pair Insulated from Human IRE1 Activity

So far, we demonstrated that cytosolic mRNA splicing could be used as an inducible post-transcriptional mechanism to restore the correct reading frame of an engineered output precursor mRNA, resulting in a protein coding mRNA that is translated into the desired output protein. However, in wild-type cells, the endogenous IRE1 activity could also process the engineered output precursor, thus limiting the functionality to IRE1 knockout cells. To make the system broadly applicable, it was crucial to identify an orthogonal mRNA hairpin-RNase pair that met three key criteria: 1) the orthogonal mRNA hairpin must not be spliced by human IRE1, ensuring insulation from endogenous processing; 2) the engineered orthogonal IRE1 must efficiently splice the orthogonal hairpin; and 3) the orthogonal IRE1 must not process the endogenous XBP1 mRNA, thus preventing unintended activation of the unfolded protein response. To this end, we examined the cytosolic kinase-RNase domains of IRE1 homologs from evolutionarily distant species, such as yeast, plant, fungi with documented cytosolic splicing^51–55^, using sequence alignments and AlphaFold3 structural predictions (Supp. Fig. 2a-c). All the selected RNase homologs exhibited substantial amino acid divergence from the predicted human RNase residues interacting with the RNA hairpin (Supp. Fig. 2d), suggesting that these homologs may possess distinct substrate specificity. To ensure diversity in their RNA cleavage site, we further predicted the corresponding XBP1 hairpin secondary structures of each species using mFold. This analysis confirmed distinct bifurcated stem-loop folding patterns with key positional differences relative to the human sequence (Supp. Fig. 3a, b), suggesting that the selected IRE1 homologs may recognize distinct RNA substrates compared to the human IRE1. For each orthologous hairpin, we inserted the bifurcated stem-loop upstream of the out-of-frame fluorescent mScarlet. Finally, to ensure the correct folding of each hairpin outside of its natural context, we flanked the hairpin sides with two 15-nt complementary regions (Supp. Table S1). To quantify the splicing efficiency of the orthologous mRNA hairpins by human IRE1, we transfected them into HEK293 cells and induced the endogenous unfolded protein response pathway with tunicamycin. We observed a notable decline in splicing processing for each species reporter, with induced fluorescent levels 10- to 100-fold lower compared to those achieved with the human IRE1 hairpin (Fig. 3a). Having validated the orthogonality of the exogenous hairpins with respect to human IRE1, we then codon-optimized their cognate kinase-RNase domains from their respective species for human expression and fused them to the FKBP^F36V^ variant^56^, which homodimerizes in response to the small molecule B/B. To systematically evaluate both the splicing efficiency and insulation of all IRE1 RNase-hairpin pairs from the endogenous human pathway, we transfected each FKBP-fused IRE1 variant, including the human, alongside each candidate mRNA hairpin, generating all possible pairwise combinations. To prevent interference from endogenous IRE1 activity, we conducted the experiment in IRE1-knockout HeLa cells, ensuring that observed splicing was solely due to the transfected constructs. Fluorescent output was then measured after B/B addition to assess splicing efficiency across conditions. Finally, we used this measured fluorescence to calculate an insulation score for each mRNA hairpin-RNase pair using three metrics (see Methods): 1) the fold reduction in splicing processing by the endogenous human IRE1 towards an orthologous hairpin, 2) the hairpin splicing efficiency ratio between the exogenous IRE1 variant and the human IRE1, and 3) the fluorescent level achieved by an exogenous IRE1-exogenous hairpin pair compared to the human IRE1-human hairpin pair. This insulation score quantifies how effectively a given IRE1 species can splice a target hairpin mRNA without interference from the endogenous IRE1 pathway.

**Figure 3.**
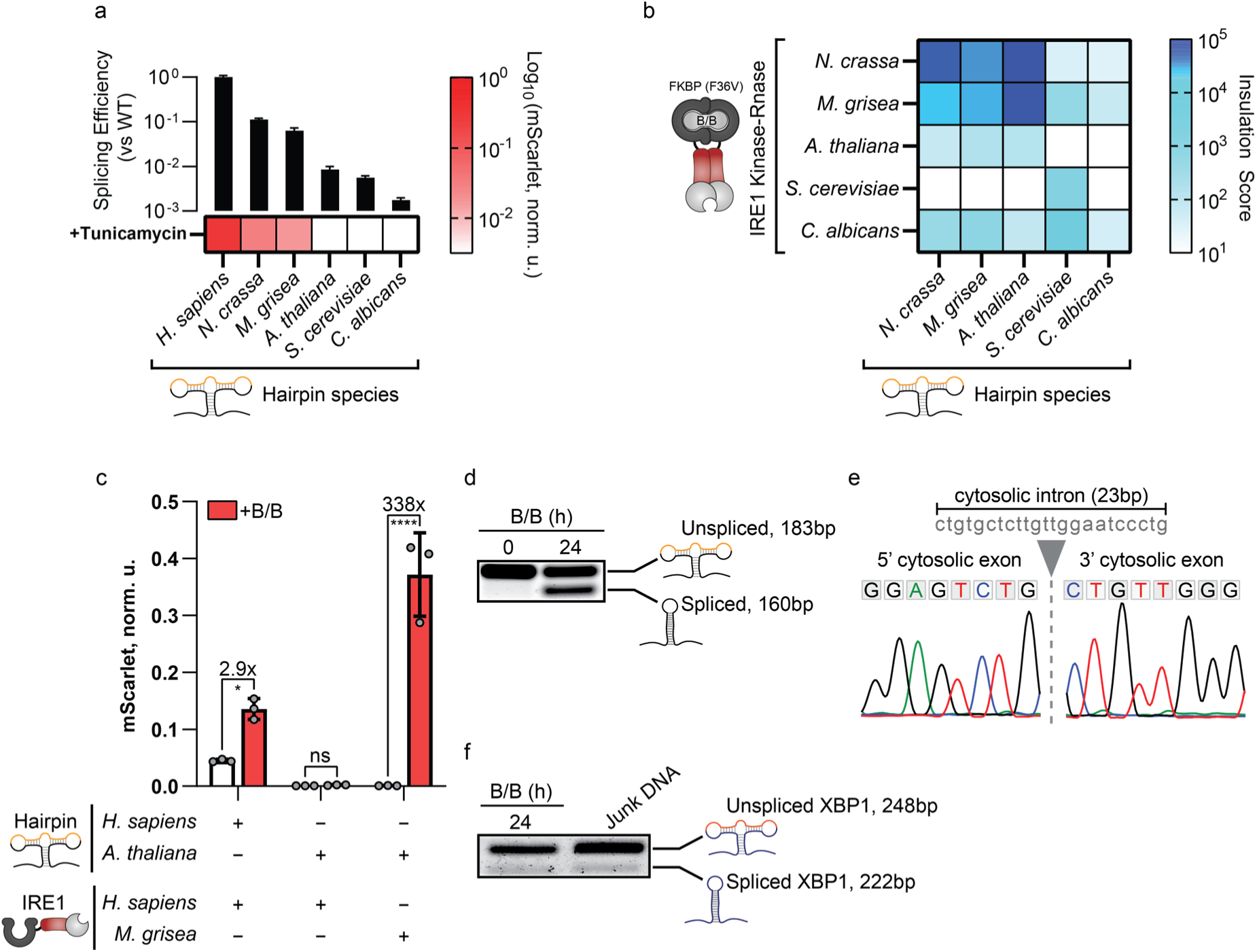
Development of SMS: an Orthologous IRE1-Hairpin Pair Insulated from Human IRE1 Activity. **a,** Human IRE1 splicing activity on different orthologous XBP1-like hairpins in HEK293 cells activated with tunicamycin. The heatmap shows the log_10_-transformed mScarlet normalized units (norm. u.) for biological triplicates, n = 3. The different hairpin species are denoted at the bottom (*Homo sapiens*, *Neurospora crassa*, *Magnaporthe grisea*, *Arabidopsis thaliana*, *Saccharomyces cerevisiae*, and *Candida albicans*), and bar charts on top depict the mean fold change relative to the human wild-type (WT) hairpin sequence, with error bars indicating the standard deviation. **b,** Heatmap of the pairwise comparison matrix of insulation scores for various XBP1-like hairpins (bottom) and IRE1-like splicing domains (left) fused to the B/B homodimerized FKBP^F36V^. Higher scores indicate better insulation from human IRE1 activity. Data are the log_10_-transformed insulation score mean for biological triplicates, n = 3. **c,** mScarlet normalized units (norm. u.) of HEK293 cells expressing the top-scoring insulated pairs of the splicing reporter (*A. thaliana*) and FKBP^F36V^-IRE1 domains (*M. grisea*) or the human components. Each dot represents an individual data point of the biological triplicates (n = 3), with error bars indicating the standard deviation. ANOVA with Šidák’s correction was performed to compare the mean of each treated column with its respective control, and asterisks indicate p < 0.05 (*) or p < 0.0001 (****). **d,** Semi-quantitative RT-PCR gel showing the unspliced (top, 183 base pairs) and spliced (bottom, 160 base pairs) forms of the *A. thaliana* hairpin in the presence of FKBP^F36V^ fused with the IRE1 domain of *M. grisea*. The appearance of the spliced band occurs upon FKBP^F36V^ homodimerization with the B/B homodimerizer. **e,** Sanger sequencing alignment of the *A. thaliana* spliced band with the predicted exon-exon boundary of the splicing reporter. The excision point of the 23-nucleotide cytosolic intron (in gray) is shown with a vertical dashed line between the 5’ and the 3’ cytosolic exons. The chromatogram colors for the nucleotides are as follows: Adenine (A in green), Thymine (T, in red), Guanine (G, in black), and Cytosine (C, in blue). **f**, Semi-quantitative RT-PCR gel showing the unspliced (top, 248 base pairs) and spliced (bottom, 222 base pairs) forms of the endogenous human XBP1 mRNA in HEK293 cells transfected with either DNA encoding FKBP^F36V^ fused with the IRE1 domain of M. grisea and treated with B/B for 24 h (left) or junk DNA (right). No increase in the spliced XBP1 band is observed upon SMS activation.

The matrix of the insulation scores reveals that several pairs are able to work efficiently, while being orthogonal with respect to human IRE1 activity (Fig. 3b, Supp. Fig. 4a). Surprisingly, cross-species activity not only occurs but also exceeds intraspecies processing for certain variants. This can be attributed to the generally higher catalytic activity of some IRE1 homologs (Supp. Fig. 4b), possibly influenced by enhanced mammalian expression due to the codon usage choice or increased solubility of their cytosolic domains compared to the corresponding intraspecies IRE1-hairpin pairs. We then selected the pair with the top-performing insulation score, comprising the plant *A. thaliana* hairpin with the fungal *M. grisea* IRE1, to validate its performance in wild-type HEK293 cells where endogenous IRE1 is naturally expressed. To this end, we transfected these cells with either the *A. thaliana* hairpin–*M. grisea* IRE1 fusion to FKBP^F36V^ or the human hairpin-human IRE1 fusion to FKBP^F36V^, and measured fluorescence signal from each splicing reporter prior to and following B/B addition (Fig. 3c). In the absence of B/B, the human hairpin-IRE1 pair exhibited strong fluorescence above background levels, indicating that endogenous IRE1 actively spliced the human hairpin in the reporter construct. In contrast, the *A. thaliana* hairpin–*M. grisea* IRE1 pair remained completely inactive, demonstrating not only that the orthogonal hairpin was unaffected by native IRE1 but also that *M. grisea* IRE1 does not splice the reporter construct in the absence of its input ligand. Upon B/B addition, the orthogonal pair exhibited strong activation achieving a dynamic range of over 300-fold, far surpassing the 2.5-fold observed with the fully human system. RT-PCR on the total RNA of transfected cells confirmed the appearance of the reporter spliced band solely after B/B treatment (Fig. 3d), while Sanger sequencing further verified the excision of the previously described^54^ 23 bp *A. thaliana* cytosolic intron (Fig. 3e). Finally, RT-PCR also confirmed that *M. grisea IRE1* does not splice the endogenous XBP1 transcript upon B/B-induced homodimerization, demonstrating that the engineered system operates without interfering with the native IRE1 pathway (Fig. 3f).

Therefore, by combining the *A. thaliana* mRNA hairpin and *M. grisea* IRE1 cytosolic fragment, we generated an orthogonal signaling unit within human cells, which we call the **s**ynthetic **m**RNA **s**plicing (SMS) system.

### SMS Receptors Enable Soluble and Surface Antigen Detection via mRNA-only Delivery

Since the SMS system does not require DNA components, it is expected to function as an mRNA-only device. To prove this hypothesis, we produced two mRNA molecules that encoded the homodimerizable FKBP^F36V^-SMS and the splicing reporter, using the T7 in-vitro transcription (IVT) system (Fig. 4a). After transient delivery to mammalian cells, the receptor mRNA is expected to be immediately translated into protein, while the splicing reporter is translated into a functionally inactive form, awaiting splicing upon the detection of the B/B molecule (Fig. 4a). Strong fluorescent signal was observed after B/B activation of the mRNA-delivered SMS sensor (Fig. 4b), with expression levels and dynamic range exceeding those of the DNA-encoded SMS pair (Fig. 4c). We then investigated whether SMS could also function as a plasma membrane mRNA-only receptor to sense extracellular signals (Fig. 4d). To this end, we designed a transmembrane version of the SMS receptor to enable extracellular signal detection. We N-terminally fused the cell surface CD4 signal peptide to FKBP^F36V^ to force surface localization, and connected it to the intracellular SMS splicing domains via the Glycophorin A (GpA) transmembrane domain, which has already proven effective in other dimerizing synthetic receptor systems^57^. This configuration allows for ligand-induced dimerization of the extracellular localized FKBP^F36V^ to be efficiently transmitted across the membrane, inducing dimerization of the intracellular *M. grisea* IRE1, thus triggering cytosolic splicing. We obtained over 60-fold induction of the mRNA-delivered SMS receptor upon B/B addition to cell culture media, with a dose-response curve showing an EC50 of 1.1 nM (Fig. 4e). However, we also noticed a slightly higher off-state signal compared to the fully intracellular version of this SMS, probably due to ligand-independent dimerization caused by membrane crowding.

**Figure 4.**
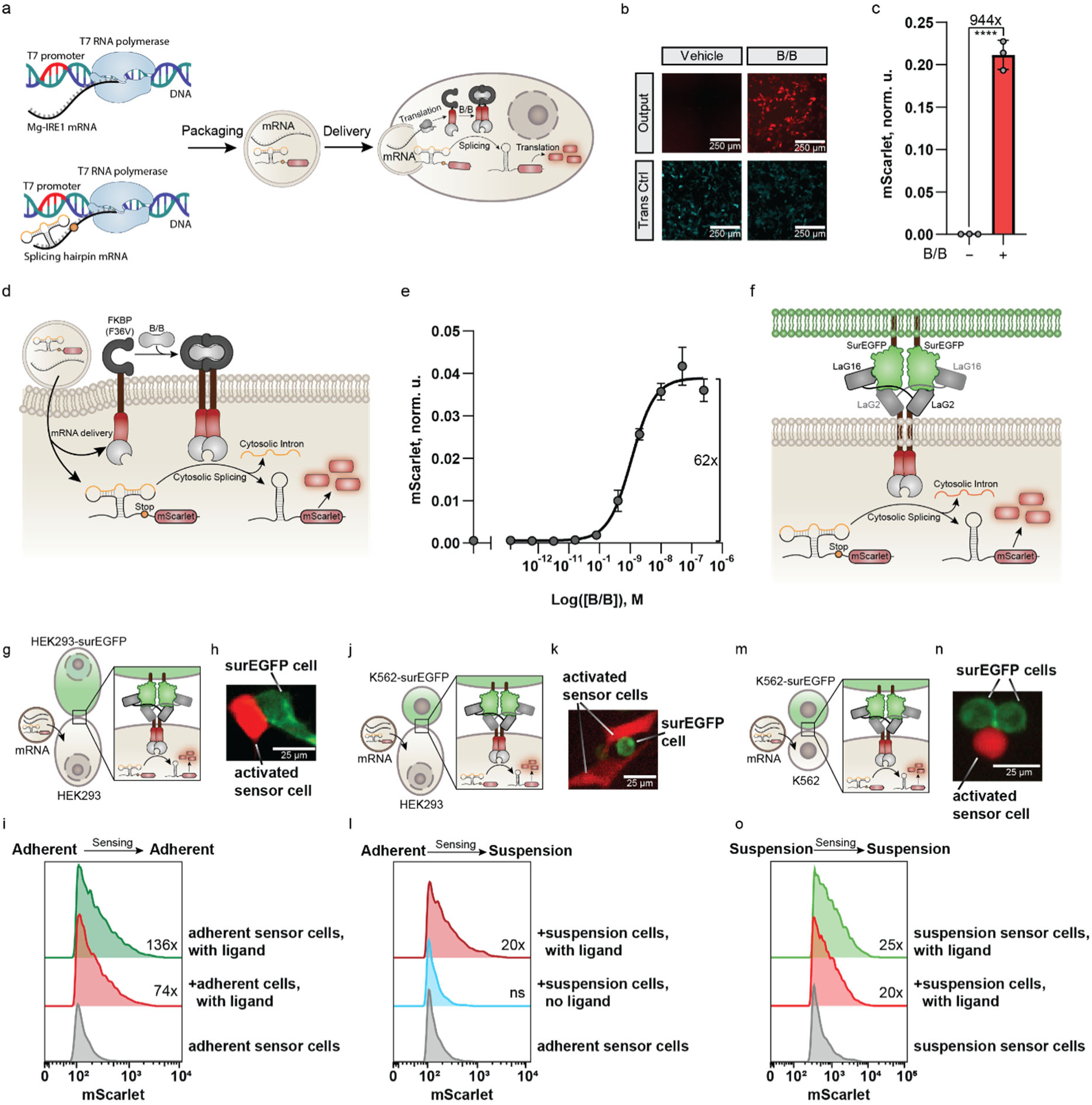
SMS Receptors Enable Detection of Soluble and Surface Antigens with mRNA-Exclusive Encoding. **a,** Schematic illustrating the production, delivery, and execution of the SMS receptor system with mRNA-only components. The PCR-linearized DNA of the *M. grisea* (Mg) IRE1 and the *A. thaliana* splicing hairpin target, each containing a 120-nucleotide 3’ poly-A tail, are in vitro transcribed to mRNA with the T7 polymerase. Upon delivery, the IRE1 mRNA is translated into its protein form, ready for activation by the chosen ligand. Upon ligand-induced dimerization, the mRNA of the synthetic splicing target is spliced to its active form, allowing the translation of the output. **b,c,** Microscopy images (b) and flow cytometry quantification (c) of HEK293 cells transfected with FKBP^F36V^-SMS mRNAs and treated with B/B to induce mRNA hairpin splicing and mScarlet production. Cells were co-transfected with control mRNA producing constitutive mCerulean for normalization. Scale bars represent 250 μm. The bar chart shows normalized mScarlet units (norm. u.) for biological triplicates (n = 3). Gray points denote individual values, with error bars indicating the standard deviation. An unpaired two-tailed t-test was performed, and asterisks indicate p < 0.0001 (****). **d,e,** Schematic showing the transmembrane FKBP^F36V^-SMS receptor (d) and its dose-response curve (e). Upon mRNA delivery, the SMS receptor is translated and localized to the plasma membrane due to the CD4 signal peptide and the glycophorin A (GpA) transmembrane domain, where it is activated by ligand-induced homodimerization resulting in activation of the synthetic splicing target. The dose-response curve shows the mScarlet fluorescence in normalized units (norm. u.) at various concentrations of the B/B homodimerizer. Each data point represents the average of biological triplicates (n = 3), with error bars indicating the standard deviation. **f,** Schematics showing the ligand-induced dimerization of the transmembrane SMS receptor upon presentation of plasma membrane-immobilized surface EGFP (surEGFP). Each nanobody subunit (Lag16 and LaG2) binds to a distinct epitope on separate surEGFP monomers, indirectly causing SMS receptor dimerization. Schematics **(g, j, m)**, microscopy images **(h, k, n),** and quantification **(i, l, o)** of membrane sensing after mRNA delivery of Lag16-LaG2-SMS components with different cell configurations: (g, h, i) adherent^surEGFP^-adherent^SMS^, (j, k, l) suspension^surEGFP^-adherent^SMS^, and (m, n, o) suspension^surEGFP^-suspension^SMS^. Scale bars represent 25 μm. Histograms show mScarlet expression in SMS-transfected populations, with each overlay representing one of the biological triplicates (n = 3).

Next, we sought to expand the range of possible extracellular protein inputs, including cell surface antigens, by incorporating nanobodies^58^ as a recognition module. To achieve that, we designed SMS receptors with an extracellular tandem fusion of two nanobodies, each recognizing a distinct epitope on the same target protein, thereby inducing dimerization upon target protein binding. We selected two nanobodies, LaG16 and LaG2, known to bind different eGFP epitopes^59^, and fused them as sensing domains of the SMS receptor to enable detection of cell-surface expressed eGFP on both adherent and suspension cell lines (Fig. 4f). We found that the mRNA-delivered LaG16-LaG2-SMS is robustly activated by eGFP recognition during cell-cell contact across adherent, suspension, and mixed adherent-suspension cell co-cultures (Fig. 4g-o). Notably, we observed that the synthetic receptor response displays a stronger signal when co-expressed with surface eGFP by the same cell (Fig. 4i, 4o), suggesting that SMS is also capable of detecting cis-presented ligand antigens. Therefore, using tandem nanobody domains as the extracellular recognition module of SMS is a programmable tool for sensing extracellular cell surface proteins with mRNA-only encoding.

### SMS Receptors Detect Clinically Relevant Inflammation Markers via mRNA-only Delivery in cell lines and primary T-cells

Next, we demonstrated the potential of SMS to sense clinically relevant molecules^60^, such as the inflammatory cytokines interleukin-1 beta (IL-1β) and tumor necrosis factor alpha (TNF-α). For the former, we designed SMS to incorporate the ectodomains of the naturally heterodimerizing IL1β complex, consisting of the interleukin-1 receptor (IL-1R^1-336^) and its accessory protein (IL-1RAcP^1-359^) extracellular regions (Fig. 5a). For the latter, we devised an SMS with the sensor domain composed of a nanobody (VHH)^61^ targeting a single subunit of the homotrimeric TNF-α, which induces receptor trimerization after TNF-a binding (Fig. 5c). Both mRNA-encoded SMSs exhibited comparable dose-response curves with purified cytokines (Fig. 5b, 5d), displaying EC50 values of about 1 nM, which matches the previously described^61, 62^ affinity of the chosen sensing subunits. To evaluate whether SMS receptors can detect endogenous cytokines levels, we tested their response to IL-1β and TNF-α produced by THP-1 monocytes upon inflammatory stimulation (Supp. Fig. 5a). First, we differentiated THP-1 cells into macrophage-like cells and induced IL-1β secretion using inflammatory signals (Fig. 5e). Supernatants collected from these stimulated cells were applied to sensor cells expressing IL-1R- and IL-1RAcP-SMS, leading to a more than 10-fold activation, confirming the detection of secreted IL-1β (Fig. 5f). Next, we assessed whether SMS could detect TNF-α in a direct co-culture system. THP-1 monocytes were stimulated with LPS to induce TNF-α secretion and were co-cultured with eGFP-tagged responder cells expressing VHH-SMS (Fig. 5g). Analysis of eGFP-positive cells revealed a more than 10-fold activation, indicating effective TNF-α sensing in a co-culture setting (Fig. 5h). In both cases, SMS activation levels closely matched the expected response based on ELISA-quantified cytokine concentrations in the supernatant (Supp. Fig. 5b, 5c), demonstrating that SMS receptors can reliably detect cell-derived cytokine levels. Importantly, we then confirmed that the SMS receptor is a functional mRNA device also in human primary T-cells (Fig. 5i). A single mRNA nucleofection of IL-1RAcP- and IL-1R-SMS was sufficient to engineer the IL-1β response of primary T-cells towards our synthetic splicing reporter (Fig. 5l). Collectively, these findings indicate that it is possible to rapidly design different SMS receptors sensing biologically relevant inputs at the mRNA level, since the cytosolic splicing reaction occurs efficiently in both cell lines and primary cells.

**Figure 5.**
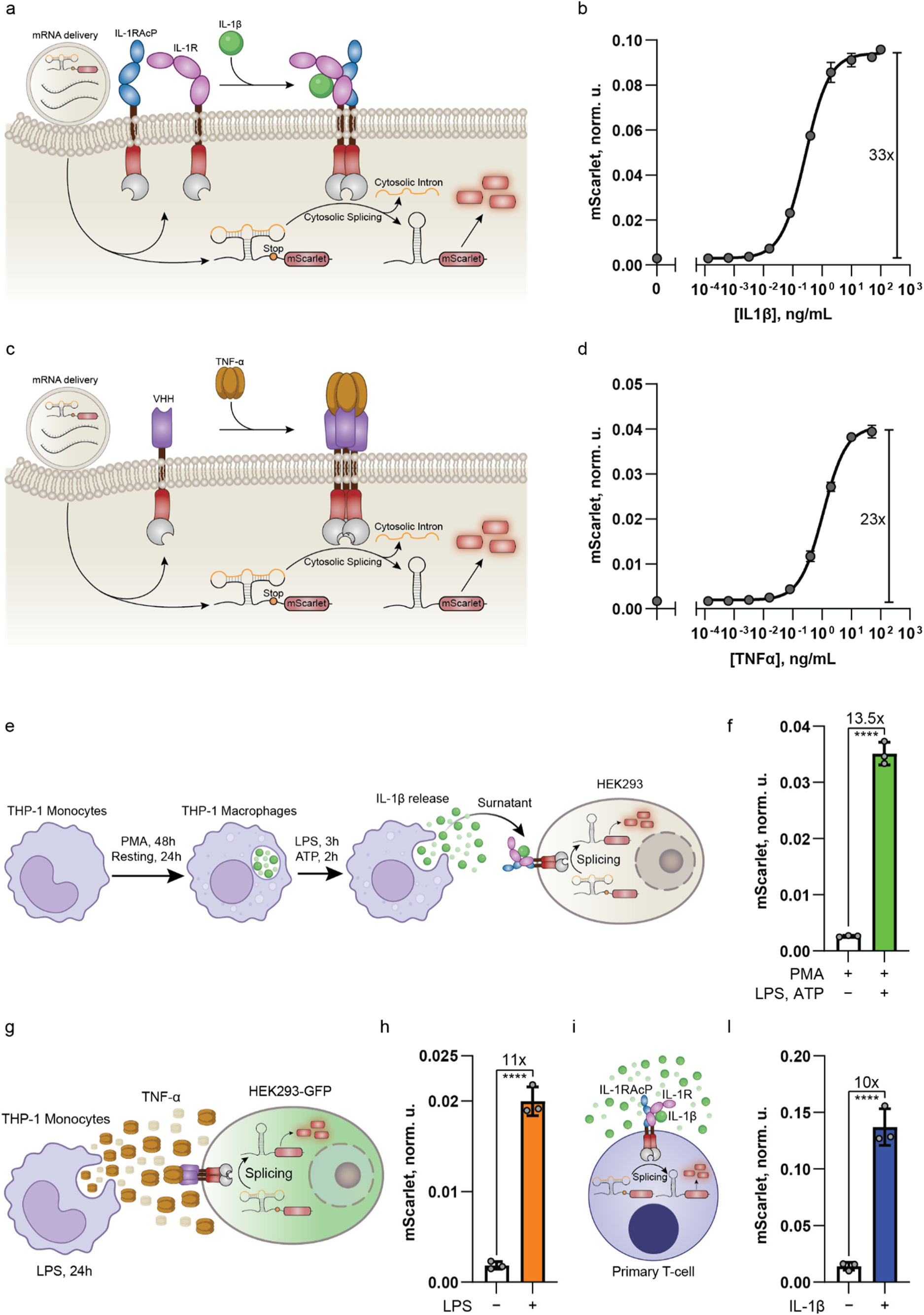
SMS Receptors Detect Biologically Relevant Inflammatory Molecules in Cell Lines and Human Primary T-cells with mRNA-Exclusive Encoding. **a,c,** Schematic illustrating the ligand-induced dimerization of the transmembrane SMS receptor after mRNA delivery. The SMS binds IL-1β through the IL-1R and IL-1RAcP ectodomains or senses TNF-α via a VHH nanobody. Upon ligand binding, the IRE1 RNase domain is activated, leading to cytosolic splicing of the synthetic mRNA reporter and enabling mScarlet translation. **b,d,** Dose-response curves showing mScarlet normalized units (norm. u.) in response to varying concentrations of IL-1β (b) or TNF-α (d). Each data point in gray represents the average of biological triplicates (n = 3), with error bars indicating the standard deviation. **e,** Diagram depicting the differentiation of THP-1 monocytes into macrophages to release IL-1β. THP-1 cells are treated with 25 nM PMA for 48 hours (h), rested for 24 h, and then stimulated with 1 μg/mL LPS for 3 h, followed by 5 mM ATP for 2 h. **f,** Quantification of splicing reporter fluorescence after detection of IL-1β-containing medium collected from differentiated THP-1 macrophages by HEK293 cells transfected with SMS mRNAs. **g,** Schematic illustrating the sensing of TNF-α release by LPS-stimulated THP-1 monocytes using co-cultured HEK293^GFP^ cells expressing the SMS receptors. **h,** Bar chart quantification of mScarlet signal in the GFP^positive^ HEK293 population, upon co-culture with THP-1 monocytes. **i,l,** Schematic illustrating the sensing mechanism and quantification of IL-1β detection by human primary T-cells delivered with SMS mRNAs. All bar charts show mScarlet normalized units (norm. u.) for biological triplicates (n = 3), with gray points denoting individual values, and error bars indicating the standard deviation. For each condition, an unpaired two-tailed t-test was performed, and asterisks indicate p < 0.0001 (****).

### Cytokine-Converter SMS Receptors transduce inflammation signals into therapeutically relevant outputs with mRNA-only encoding

To verify that the cytokine-sensing SMS receptors can produce significant levels of a desired therapeutically relevant protein, we replaced the fluorescent reporter with the out-of-frame anti-inflammatory IL-10 immunomodulator downstream of the splicing hairpin (Fig. 6a). To ensure the independent translation of the output protein from the coding sequence of the spliced hairpin, we placed a T2A element immediately in front of the IL-10 cytokine. In this way, we achieved cytokine-converter SMS receptors, in which the sensing of an inflammatory input (TNF-α or IL-1β) is transformed to the extracellular release of an anti-inflammatory output (IL-10). The mRNA administration of the cytokine-converter IL-1R- and IL-1RAcP-SMS (Fig. 6b) in HEK293 cells was able to produce high IL-10 levels in response to IL-1β, either secreted by stimulated THP-1 macrophages or via direct addition to the medium (Fig. 6c). Likewise, LPS-induced TNF-α release was efficiently detected by sensor cells equipped with the VHH-SMS mRNA (Fig. 6d), with output IL-10 amounts comparable to those obtained with the IL-1β receptor variants (Fig. 6e). Notably, we also found that the dynamic range was minimally affected by a ∼10-fold increase in the transfection amounts of the splicing hairpin mRNA (Supp. Fig. 6a, c), while IL-10 production increased almost linearly (Supp. Fig. 6b, d). This result suggests that the output levels can be easily fine-tuned to the desired therapeutic concentration by simply dosing of the splicing hairpin mRNA. Remarkably, when we tested the IL-1β cytokine-converter SMS in primary T-cells by mRNA nucleofection (Fig. 6f), we measured IL-10 production equivalent to that obtained in cell lines (Fig. 6g), which we confirmed to not occur naturally upon IL-1β stimulation alone (Supp. Fig. 6e, f). These results further highlight that SMS is a modular mRNA-only device, providing programmable and tunable input-output mRNA splicing reactions in human primary cells and supporting a clinically-relevant sense-response behavior.

**Figure 6.**
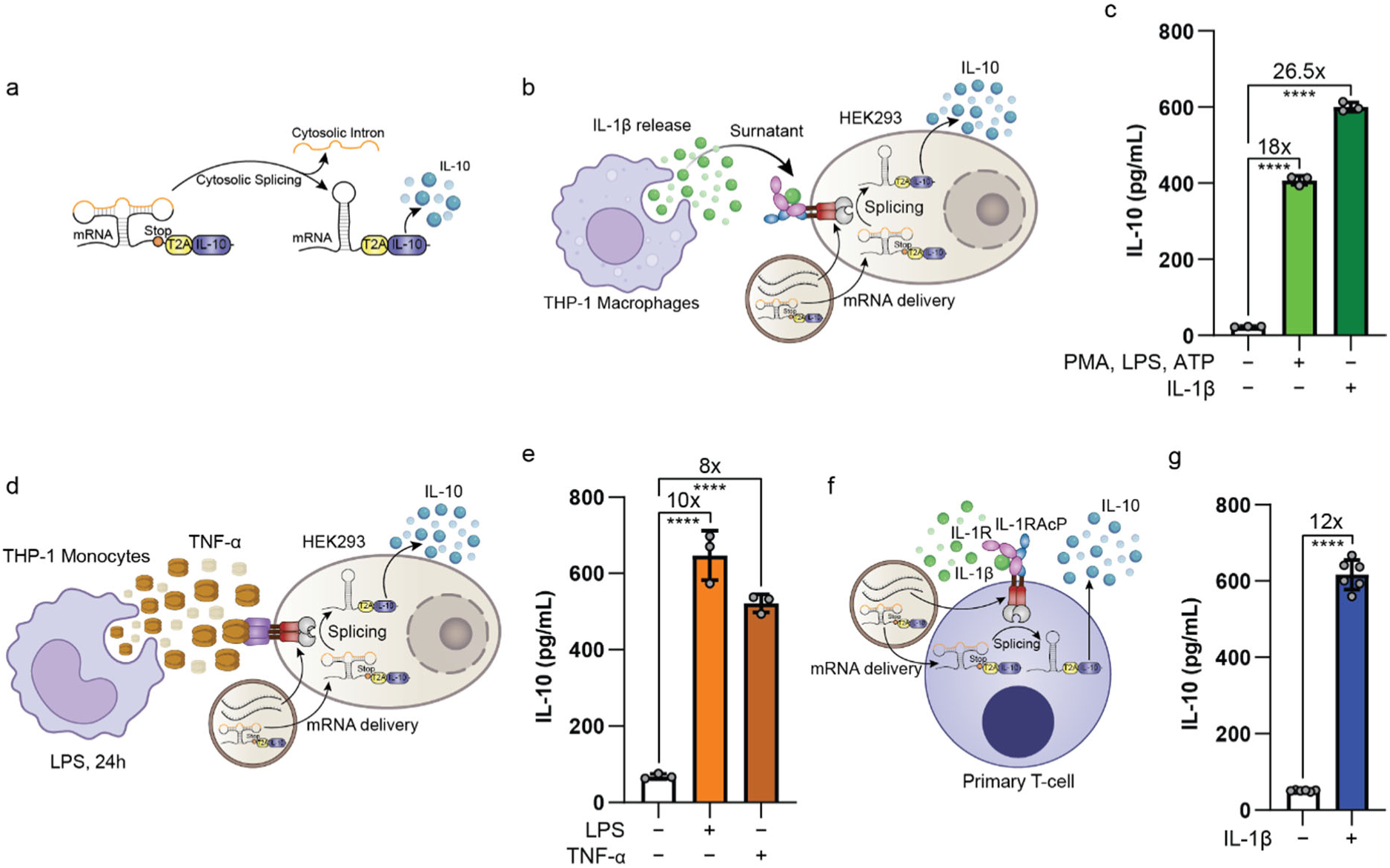
SMS Receptors Secrete the Anti-inflammatory IL-10 cytokine Following Inflammatory Detection in Cell Lines and Human Primary T-cells with mRNA-Exclusive Encoding. **a,** Design of the IL-10 synthetic mRNA splicing output. The mRNA encoding the IL-10 cytokine is placed out-of-frame downstream of the *M. grisea* splicing hairpin and a T2A self-cleaving peptide sequence. Upon cytosolic intron removal, the T2A ensures that the IL-10 cytokine is translated into the secretory pathway with its native sequence. **b,c,** Schematic illustrating the sensing of IL-1β released by differentiated THP-1 macrophages and ELISA quantification of IL-10 production by mRNA-delivered HEK293 cells. ANOVA with Dunnett correction was performed to compare each condition to the control column (n = 3). **d,e,** Schematic illustrating the sensing of TNF-α released by THP-1 monocytes and ELISA quantification of IL-10 production by mRNA-delivered, co-cultured HEK293 cells. ANOVA with Dunnett correction was performed to compare each condition to the control column (n = 3). **f,g,** Schematic illustrating the delivery of SMS mRNAs into human primary T-cells. Detection of IL-1β through IL-1R and IL1-RAcP leads to the activation of the IRE1 RNase domains, causing the cytosolic splicing of the IL-10 synthetic mRNA and the subsequent extracellular release of the mature cytokine protein product, quantified by ELISA. An unpaired two-tailed t-test was used for statistical analysis (n = 6). All bar charts show single data points in gray, with error bars representing the standard deviation and asterisks indicate p < 0.0001 (****).

## Discussion

Implementing customizable sensory responses with mRNA-encoded components has been a challenging goal in synthetic biology and a limiting factor for programmable mRNA-based therapy. The key missing element has been a modular and insulated platform capable to transmit information post-transcriptionally in response to desired molecular cues. Here, we show that orthologous components of the mRNA cytosolic splicing pathway fulfill the stringent requirements for such a platform, being able to convert extracellular or intracellular stimuli into the translational activation of an arbitrary engineered mRNA target to generate a protein of interest. The SMS receptors demonstrate remarkable versatility in sensing diverse inputs, including small molecule ligands, cell-bound antigens, and biologically relevant extracellular soluble proteins, through mRNA-based encoding, and are functional in both common cell lines and human primary cells. The sensor region can be constructed not only with naturally dimerizing domains but also with tandem single-domain antibody fusions, providing access to a plethora of already available targets. On the output side, virtually any protein of interest can be rapidly placed under the control of SMS by incorporating a small RNA hairpin region upstream of its coding sequence. In addition, the SMS system retains the general features of the unfolded protein response pathway. It is a multiple-turnover reaction, in which each active receptor catalytically processes several mRNA molecules^63^. This enzymatic mechanism ensures signal amplification and avoids the formation of signaling-dead receptors^16^, a prevalent issue in most previous DNA-based synthetic receptors. To our knowledge, this is the first time that mRNA cytosolic splicing has been engineered for synthetic modulation, leaving several aspects to be further explored and optimized in the future. For instance, using diverse transmembrane domains could modulate receptor performance to reach the hundred-fold induction seen with the intracellular SMS, because the choice of membrane region can significantly affect receptor activity^64^. Although in this work we mainly explored mRNA delivery, our findings indicate that SMS is equally suitable for DNA encoding. This extends its use to the broad range of well-established genetically encoded applications, such as genomic stable integration for long-lasting activity, while preserving the advantage of being a transcription-factor independent system. Overall, we envision that SMS receptors will prove to be a relevant tool in the synthetic engineering of mammalian cells, with pivotal importance in advancing off-the-shelf mRNA medicine to the next phase.

## Methods

### Recombinant DNA methods

Plasmids were generated using standard cloning techniques (See Table S1 for detailed sequences), and all kits were used without deviation from the manufacturer’s protocols. Restriction enzymes were purchased from New England Biolabs (NEB) or Thermo Fisher Scientific. Oligonucleotides used as primers or for oligo cloning were ordered as desalted or PAGE-purified from Integrated DNA Technologies (IDT) or Sigma-Aldrich. Polymerase Chain Reaction (PCR) was performed using Phusion High-Fidelity DNA Polymerase (NEB; cat# M0530) or the Quick-Load® Taq 2X Master Mix (NEB; cat# M0271L). PCR products and digestion fragments were gel-purified using the QIAquick Gel Extraction Kit (Qiagen; cat# 28706) or the QIAquick PCR Purification Kit (Qiagen; cat# 28106), based on the expected size of the DNA band. Ligations were performed using T4 DNA Ligase (NEB; cat# M0202) at 4 °C overnight. Synthetic sequences longer than 300 bp were ordered as DNA fragments from Twist Bioscience and assembled using Gibson assembly. The Gibson assembly reaction was carried out at 50°C for 1 hour in a 20 μL final volume, with the vector (100 ng) and inserts (3-5 molar equivalents) combined in 1x Gibson Master Mix. Cloned plasmids were transformed into chemically competent E. coli, using TOP10 cells for non-lentiviral plasmids and STBL4 cells (ThermoFisher; cat# C737303) for lentiviral plasmids. Bacteria were plated on LB agar plates containing 100 μg/mL ampicillin (Sigma-Aldrich; cat# A9518) as a selection agent. Correct clones were identified through single E. coli colony sequencing performed by Microsynth AG, and further verified by full-plasmid sequencing using nanopore technology. Plasmid isolation was performed using PureYield Plasmid Midiprep System (Promega; cat#A2495), followed by a further purification with the Endotoxin Removal Kit (Norgen; cat# 52200).

### Mammalian cell culture

HEK293 cells (Life Technology; cat# 11631-017) and HEK293T cells (ATCC; cat# CRL-11268) were maintained in DMEM medium (Gibco; cat# 41966-029). HeLa IRE1 knockout cells carrying a 1 bp deletion in exon 10, a 1 bp insertion in exon 10, and a 2 bp deletion in exon 10 generated with CRISPR/Cas9 were purchased from ATCC (ab255389) and also cultured in DMEM medium. K-562 cells (ATCC; cat# CCL-243) and THP-1 monocytes (ATCC; cat# TIB-202) were grown in RPMI 1640 medium (Gibco; cat# 72400-021). Purified primary T-cells were maintained in RPMI 1640 medium supplemented with 200U/mL of interleukin-2 (STEMCELL Technologies; cat# 78220). All cell cultures were maintained at 37°C in a humidified atmosphere with 5% CO2, using the described culture medium supplemented with 10% fetal bovine serum (FBS) (Gibco; cat# 10270-106) and 1% penicillin-streptomycin solution (P/S) (Corning; cat# 30-002-CI), hereafter referred to as the complete medium. During co-culture experiments, cells were kept in complete DMEM medium. Mycoplasma contamination testing was conducted using the PCR mycoplasma test kit (Promokine; cat# PK-CA91-1024), following the manufacturer’s instructions.

### THP-1 Macrophage differentiation for IL-1β secretion stimulation

1.0 x 10^6^ THP-1 monocytes (ATCC; cat# TIB-202) were seeded in 24-well plates in 500 μL of medium and differentiated into macrophages by incubation of 25 nM phorbol 12-myristate 13-acetate (PMA) (Abcam; cat# ab147465) for 48 hours. Following this incubation, the cells underwent a 24-hour resting period during which the medium was replaced to remove the PMA. The next day, IL-1β secretion was induced by stimulation with 1 μg/mL Escherichia coli lipopolysaccharide (LPS) (MERCK; cat# L5543) for 3 hours, followed by the addition of 5 mM adenosine triphosphate (ATP) (Invivogen; cat# tlrl-atpl) for 2 hours. The medium was then collected to measure the quantity of released IL-1β or immediately used for further experiments.

### Generation of stably expressing surface eGFP cell lines

Lentivirus was produced by co-transfecting the packaging plasmids (10 μg of pCMV-Gag-Pol, 2 μg of pCMV-VSVG, and 1 μg of pRSV-Rev) and the transfer plasmid (15 μg of pEF1a-surface eGFP) into 3.8 x 10^6^ HEK293T cells seeded on a 60.1 cm² dish using a ratio of Polyethylenimine ‘Max’ (Polysciences; cat# 24765) (PEI) 3 to 1 (DNA) in a total volume of 1mL of Opti-MEM. Eighteen hours post-transfection, the media was exchanged with 15 mL of complete medium. Two media collections were performed at 48- and 72-hours post-transfection, pooled, centrifuged and sterile filtered (0.45 μm) (Sartorius; cat# 16555-K). The supernatant was then concentrated using Amicon Ultra-15 Centrifugal Filter Unit (Merck Millipore; cat# UFC9100) by centrifugation and buffer exchange was performed with 10mL of PBS-MK (1 mM MgCl_2_, 2.5 mM KCl in PBS), concentrated to a final volume of ∼1.2 mL and stored at -80°C. The transduction was performed by adding 200 μL of the concentrated lentiviral solution to HEK293 or K-562 cells seeded in 2 mL of complete medium in a 6-wells (Thermo Scientific; cat# 140675) plates at a density of 2.4 x 10^5^ and 4.0 x 10^5^, respectively. Sorting was performed 5 days after transduction to obtain a pure population of eGFP expressing cells using the BD FACSAria™ Fusion (Ex: 488 nm, bandpass filter 530/30 nm, longpass filter 502 nm).

### Human Primary T-cells isolation

Peripheral Blood Mononuclear Cells (PBMCs) were isolated from fresh buffy coats. Buffy coats were diluted 1:1 in PBS containing 2% FBS and 2 mM EDTA, layered over 10 mL of Ficoll gradient, and centrifuged at 400g for 40 minutes. The mononuclear cell layer was collected, washed twice with PBS containing 2% FBS and 2 mM EDTA, and red blood cells were lysed using RBC lysis buffer (Invitrogen™ cat# 00-4333-57). Cells were then cryopreserved using Bambanker™ DIRECT (Nippon Genetics, cat# BBD01) and stored in liquid nitrogen. For isolating T-cells, PBMCs were thawed in a 37°C water bath, and washed with pre-warmed complete RPMI medium. Cells were then resuspended in FACs buffer (PBS, 2% FBS, 1 mM EDTA) and incubated with DNase I Solution (100 μg/mL) (STEMCELL Technologies cat# 07900) at room temperature for 15 minutes. Aggregated suspensions were filtered through a 37 μm cell strainer, and cells were resuspended in FACs buffer at 5 x 10^7^ cells/mL for T-cell enrichment. T-cell enrichment was performed using the EasySep™ Human T-Cell Isolation Kit (STEMCELL Technologies; cat# 17951). PBMCs were incubated with the EasySep™ Human T-Cell Isolation Cocktail and RapidSpheres™ according to the manufacturer’s instructions, followed by magnetic separation with the EasyEights™ EasySep™ Magnet (STEMCELL Technologies; cat# 18103).

### mRNA synthesis

mRNA synthesis was carried out using the Invitrogen™ mMESSAGE mMACHINE™ T7 Transcription Kit (Invitrogen™; cat# AM1344). Plasmid DNAs were linearized by PCR using a constant forward and reverse primer pair that binds to common 5’ and 3’ regions shared across all constructs. The forward primer included the T7 promoter recognition sequence (5’-TAATACGACTCACTATAGGGAGA-3’), while the reverse primer introduced a 120nt poly-A tail. The linearized PCR products were then gel-purified with the QIAquick Gel Extraction Kit (Qiagen; cat# 28706) and normalized to 100 ng/μL. The in vitro transcription reaction was conducted according to the manufacturer’s protocol, utilizing 2 μL of the normalized DNA as input. The reaction mixture was incubated at 37°C for 2 hours to ensure efficient mRNA synthesis, followed by a 15-minute DNase I treatment at 37°C to remove the DNA template, then purified using the MEGAclear Transcription Clean-Up Kit (Invitrogen™; cat# AM1908).

### mRNA Delivery

mRNA delivery in HEK293 and K-562 cells was performed with Lipofectamine™ MessengerMAX™ (Invitrogen™; cat# LMRNA008). HEK293 were seeded in 24-well plates (Thermo Scientific; cat# 142475) at a density of 1.5 x 10^5^ in complete DMEM and transfected the following day. K-562 cells were seeded were seeded on 24-well plates at a density of 2 x 10^5^ in complete RPMI and transfected on the same day. For transfection, the in vitro transcribed mRNAs were combined and diluted Opti-MEM (Gibco; cat# 31985062) to a final volume of 50 μL per transfection. Separately, Lipofectamine™ MessengerMAX™ was diluted in Opti-MEM, incubated for 10 min at RT, and added to the mRNA-Opti-MEM mix to a total volume of 100 μL with a Lipofectamine™ MessengerMAX™ (μL) to mRNA (μg) ratio of 2:1. After a 5-minute incubation at RT, the transfection mix was added dropwise to the cells. Primary T-cells were activated with Dynabeads™ Human T-Activator CD3/CD28 (Gibco cat# 11131D) 48 hours prior to mRNA delivery by nucleofection. Briefly, T-cells were seeded at 8 x 10^4^ in 200 μL of medium in a 96-well plate (PerkinElmer; cat# 6005182), and 2 μL of Dynabeads™ were added to each well. On the day of nucleofection, primary T-cells were pooled from the 96-well plate, and Dynabeads™ were removed by magnetic separation with the EasyEights™ EasySep™ Magnet (STEMCELL Technologies; cat# 18103). Nucleofection was performed using the 4D-Nucleofector® X Unit (Lonza; cat# AAF-1003X) with the P3 Primary Cell 4D-Nucleofector™ X Kit S (Lonza; cat# V4XP-3032) in 16-cuvette strips. For each cuvette, 5 x 10^5^ Primary T-cells were resuspended in 20 μL of ice-cold P3 primary cell solution. The appropriate amounts of mRNAs were then directly pipetted into the cell suspension, and the T-cell human stimulated protocol was executed. Immediately after nucleofection, 80 μL of 37°C warm medium was added to the cuvette, and the cells were incubated for 10 minutes before being transferred to 24-well plates containing 500 μL of 37°C warm medium. For co-culture experiments, mRNA delivery was first performed on the intended cell population, followed by the addition of the second cell population 4 hours later at a 1:1 ratio.

### DNA Transfection

HeLa IRE1 KO and HEK293 cells were seeded in 24-wells plate at a density of 6 x 10^4^ and 7.5 x 10^4^, respectively. The following day, HeLa IRE1 KO were transfected using Lipofectamine 3000 (L3K) (Invitrogen™; cat# L3000008) and HEK293 using Lipofectamine 2000 (L2K) (Invitrogen™; cat# 11668-027). For each L2K transfection reaction, the appropriate mix of endotoxin-free plasmid DNAs was diluted in Opti-MEM to a final volume of 50 μL. Separately, L2K was diluted in Opti-MEM and incubated for 5 minutes at room temperature. The L2K-Opti-MEM mix was then combined with the DNA-Opti-MEM mix to a total volume of 100 μL, using a L2K (μL) to DNA (μg) ratio of 2:1. Following a 15-minute incubation at room temperature, the transfection mix was added dropwise to the cells. For each L3K transfection reaction, the plasmid DNAs were diluted in Opti-MEM to a final volume of 25 μL. To this, 2 µL of p3000 reagent per µg of DNA was added and the mix was vortexed briefly, followed by a 5-minute incubation. Separately, Lipofectamine 3000 was diluted in Opti-MEM and combined with the DNA-P3000-Opti-MEM mix at L3K (μL) to DNA (μg) ratio of 1.5:1. After a 15-minute incubation at room temperature, the transfection mix was added dropwise to the cells. 16 hours post-transfection with L3K, the medium of HeLa IRE1 KO cells was exchanged with 500 µL of complete DMEM. All DNA-transfected cells were analyzed 48 hours post-transfection by microscopy and flow cytometry.

### Cell Treatments

For cell treatments, the medium was replaced with either fresh appropriate medium or medium containing the desired treatment 16 hours after DNA transfection with L3K, or 24 hours after DNA transfection with L2K. For mRNA-transfected cells, treatments were directly pipetted into the 24-well plates 2 hours after transfection. The reagents used to induce receptor sensing were A/C heterodimerizer (Takara Bio; cat# 635079), the B/B homodimerizer (Takara Bio; cat# 635059), recombinant human TNF-α (R&D cat# 210-TA-020), and recombinant human IL-1β (R&D cat# 201-LB-010). With the exception of dose-response curve studies, the concentrations used were as follows: A/C was at 1 μM, B/B was at 500 nM, and IL-1β and TNF-α at 10 ng/mL. To induce TNF-α secretion from undifferentiated THP-1 monocytes during co-culture experiments, 1 μg/mL Escherichia coli lipopolysaccharide (LPS) (MERCK; cat# L5543) was used overnight. To induce activation of the endogenous IRE1 receptor, tunicamycin (Sigma-Aldrich; SML1287) was used at 5 μg/mL.

### Cytokine measurements

All cytokine measurements were performed using the enzyme-linked immunosorbent assay (ELISA) platform (R&D Systems; cat# 600-100) with the microfluidic Simple Plex cartridge (Bio-Techne; cartridge cat# SPCKA-PS-004490) to perform an automated sandwich ELISA against the human IL-10, IL-1β, and TNF-α for each sample well. The cell medium collected for cytokine quantification was centrifuged at 1000g for 6 minutes and the resultant supernatant was diluted 1:4 by pipetting 20 μL into 60 μL of the provided sample diluent. 50 μL of the diluted sample was then pipetted into the cartridge well for the automated analysis. The cytokine amounts were then calculated using the standard curves calibrated by the manufacturer.

### Microscopy

Microscopy images were acquired on a Nikon Eclipse Ti2 inverted epifluorescence microscope equipped with a motorized stage and an environmental control chamber set to 37°C and 5% CO2. The excitation light was generated by a solid-state LED light engine and filtered through Semrock cubes. Filtered emission light was collected by a Hamamatsu ORCA-Flash4.0 camera using a 10X objective and a 500 ms exposure time. Each fluorescent protein was imaged using the following excitation (Ex) light, and emission (Em) and dichroic (Dc) filter settings: mTagBFP2 (390 nm LED with LED power at 60%, Em 483/32 nm, Dc 458 nm), mCerulean (Ex 438 nm with LED power at 60%, Em 483/32 nm, Dc 458 nm), eGFP (475 nm with LED power at 60%, Em 542/27 nm, Dc 520 nm), mScarlet (575 nm LED with LED power at 60%, Em 624/40nm, Dc 593 nm). Images were processed using Fiji v2.15.1.

### Flow cytometry

For flow cytometry analysis, HEK293 and HeLa IRE1 KO adherent cells were detached using 135 μL of a 1:1 solution of PBS (Gibco; cat# 10010-015) and phenol red-free Trypsin-EDTA (0.5%) (Gibco; cat# 15400054). Suspension K-562 and primary T-cells were directly analyzed in their cell culture medium. Cells were analyzed on a BD LSRFortessa™ Cell Analyzer (BD Bioscience), calibrated with Sphero Rainbow Calibration Particles (BD; cat# 559123). Fluorescent proteins were measured using the following excitation lasers (Ex) and emission filters (Em): mTagBFP2 (Ex: 405 nm, Em: 450/50 nm), mCerulean (Ex: 445 nm, Em: 473/10 nm), eGFP (Ex: 488 nm, Em: 530/30 nm, longpass filter 505 nm), and mScarlet (Ex: 561 nm, Em: 610/20 nm, longpass filter 600 nm). At least 100,000 cells were acquired for each biological replicate, and flow cytometry data were analyzed using FlowJo software (BD Biosciences) by first gating live cells based on forward and side scatter area, followed by gating singlets based on forward scatter height and forward scatter area. For each fluorescent protein, the positive signal threshold was determined by comparing color-positive and color-negative control cells, ensuring that no more than 0.1% of the true negative cells were mislabeled as false positives. To account for transfection variability within and across experiments, the expression value for the fluorescent signal of interest (I) was then normalized to a co-transfected internal fluorescent control (C) using the following formula:

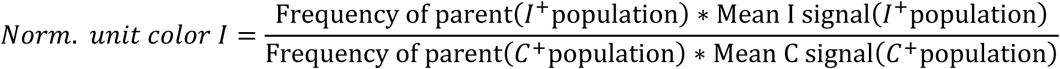

### Cytosolic mRNA Splicing Assay

Total RNA was extracted from HEK293 cells using the RNeasy Plus Mini Kit (Qiagen; cat# 74134). Cells were seeded at a density of 7.5 x 10^4^ per well in 24-well plates, and RNA was pooled from two wells for each condition. The RNA samples were resuspended in RNase/DNase-free water (Invitrogen™; cat# 10977049) and quantified by NanoDrop spectrophotometer. cDNA was synthesized by reverse transcription using the SuperScript IV First-Strand Synthesis System (Invitrogen™; cat# 18091050) with random hexamer primers, using 500 ng of total RNA as the input. The cDNA products were then diluted 1:10 with RNase-free water. PCR amplification was performed using 2 μL of the diluted cDNA with Quick-Load Taq 2x Master Mix (NEB; cat# M0271) under the following conditions: 28 cycles of denaturation at 95°C for 15 seconds, annealing at 59°C for 30 seconds, and extension at 68°C for 1 minute. A final extension at 68°C for 5 minutes. The primers used for the PCR were as follows: for both the endogenous human XBP1 and the human splicing reporter, the forward primer was GGAGTTAAGACAGCGCTTGG. The reverse primer for the endogenous human XBP1 was ACTGGGTCCAAGTTGTCCAG. The reverse primer for the human splicing reporter was TTTTGCCGTTTGCGTTCCTT. For the *A. thaliana* splicing reporter: forward primer was GCCACCATGTATCCTTATGATGT and reverse primer was TGACGGCTTCCCCTTTTGAA. The PCR products were analyzed by electrophoresis on a 2.5% agarose gel run for 3-4 hours until the bands were resolved. PCR bands were extracted using the QIAquick Gel Extraction Kit (Qiagen; cat# 28706) and sent for Sanger sequencing at Microsynth AG.

### Insulation score calculation

The insulation score for a given IRE1-Hairpin mRNA species pair quantifies how much more effectively a given IRE1 species can splice a target hairpin mRNA relative to the endogenous human IRE1 receptor, taking also into account the total levels of fluorescent protein produced by the Hairpin mRNA species compared to the human hairpin mRNA reporter. The insulation score (IS) is calculated from the normalized fluorescent values of the respective splicing reporters using the following formula:

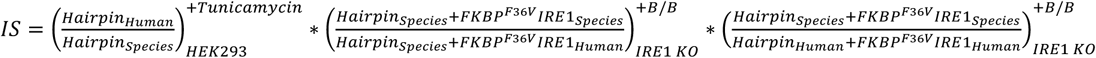

The first parenthesis represents the fold reduction in splicing processing by the endogenous human IRE1 on the orthologous hairpin activated by Tunicamycin in HEK293 cells. The second parenthesis represents the efficiency splicing ratio on the species hairpin between the chosen IRE1 species and the human IRE1 FKBP^F36V^ fusion proteins upon homodimerization by B/B in IRE1 HeLa KO cells. The third parenthesis represents the fold change of the active fluorescent level calculated between the FKBP^F36V^ fusions of the given IRE1-hairpin pair and the human IRE1-hairpin pair upon homodimerization by B/B in IRE1 HeLa KO cells.

### Data analyses

Homologous human XBP1 sequences containing the bifurcated stem loop were retrieved from the National Center for Biotechnology Information (NCBI) utilizing the Basic Local Alignment Search Tool (BLAST). Homologous human IRE1 sequences were retrieved from the Uniprot Knowledgebase (UniProtKB). RNA secondary structures were calculated with mFold on the UNAFold web server (http://www.unafold.org/) and drawings were performed using Visualization Applet for RNA (VARNA). Tertiary structures were calculated with the AlphaFold 3 web server (https://alphafoldserver.com/). Protein-Ligand Interaction Profiler (PLIP, https://plip-tool.biotec.tu-dresden.de/) was used to identify the RNA-IRE1 interactions from the predicted AlphaFold 3 structures. Multiple RNA sequence alignments were conducted with LocARNA (http://rna.informatik.uni-freiburg.de/LocARNA/), using mFold secondary structures and default parameters, with the structure weight adjusted to 300. Multiple Protein sequence alignment was performed with Clustal Omega and rendered with Easy Sequencing in PostScript (ESPript 3.0, https://espript.ibcp.fr/ESPript/ESPript/). Statistical analysis was performed using GraphPad Prism v9.2.0. To compare the means of two independent groups, an unpaired two-tailed t-test was used. When comparing the means of three or more independent groups, one-way analysis of variance (ANOVA) was applied. Post-hoc corrections included Tukey’s test, when ANOVA was used to identify significant differences between all possible pairs of groups, Dunnett’s test, when ANOVA was used to compare each group mean against the same control group mean, or Šidák’s test, when ANOVA was used to compare the means of selected pairs of columns. In all figures, bar charts and plots represent the mean values of replicate measurements, with error bars indicating ±1 standard deviation (SD). The significance threshold was set to p < 0.05, and significance levels are reported as: *p < 0.05; **p < 0.01; ***p < 0.001; ****p < 0.0001.

## Supporting information

See Table S1

---

**Supplementary Figure S1.**
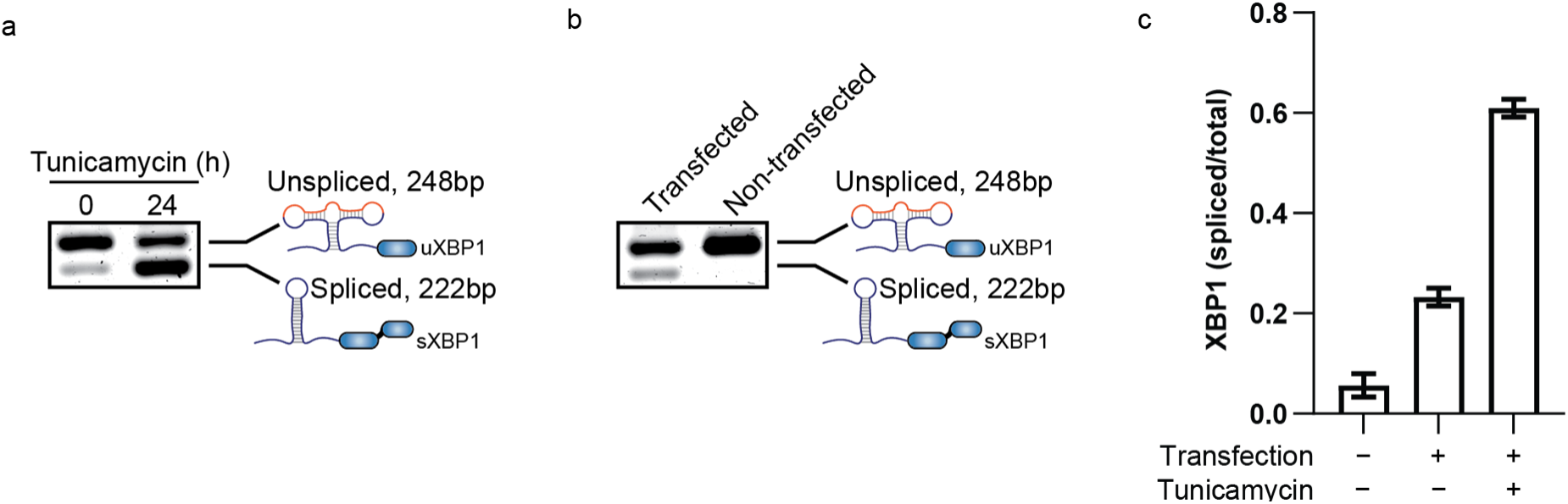
Endogenous XBP1 mRNA Splicing in HEK293 Cells. **a,** RT-PCR gel showing the unspliced (uXBP1, 248 base pairs) and spliced (sXBP1, 222 base pairs) forms of XBP1 mRNA in HEK293 cells treated with tunicamycin for 24 hours (h). **b,** RT-PCR gel depicting uXBP1 and sXBP1 bands in HEK293 cells transfected with 500 ng of junk DNA compared to non-transfected cells. **c,** Quantification of band intensities from RT-PCR gels in (a) and (b). For each condition, the intensity of the sXBP1 band was normalized to the sum of the intensities of the spliced (sXBP1) and unspliced (uXBP1) bands, representing the total XBP1 signal. The bar chart illustrates the mean spliced/total XBP1 ratio for biological duplicates (n = 2) for non-transfected cells, biological replicates (n = 9) for transfected cells, and biological triplicates for tunicamycin-treated cells (n = 3). Error bars indicate the standard deviation.

**Supplementary Figure S2.**
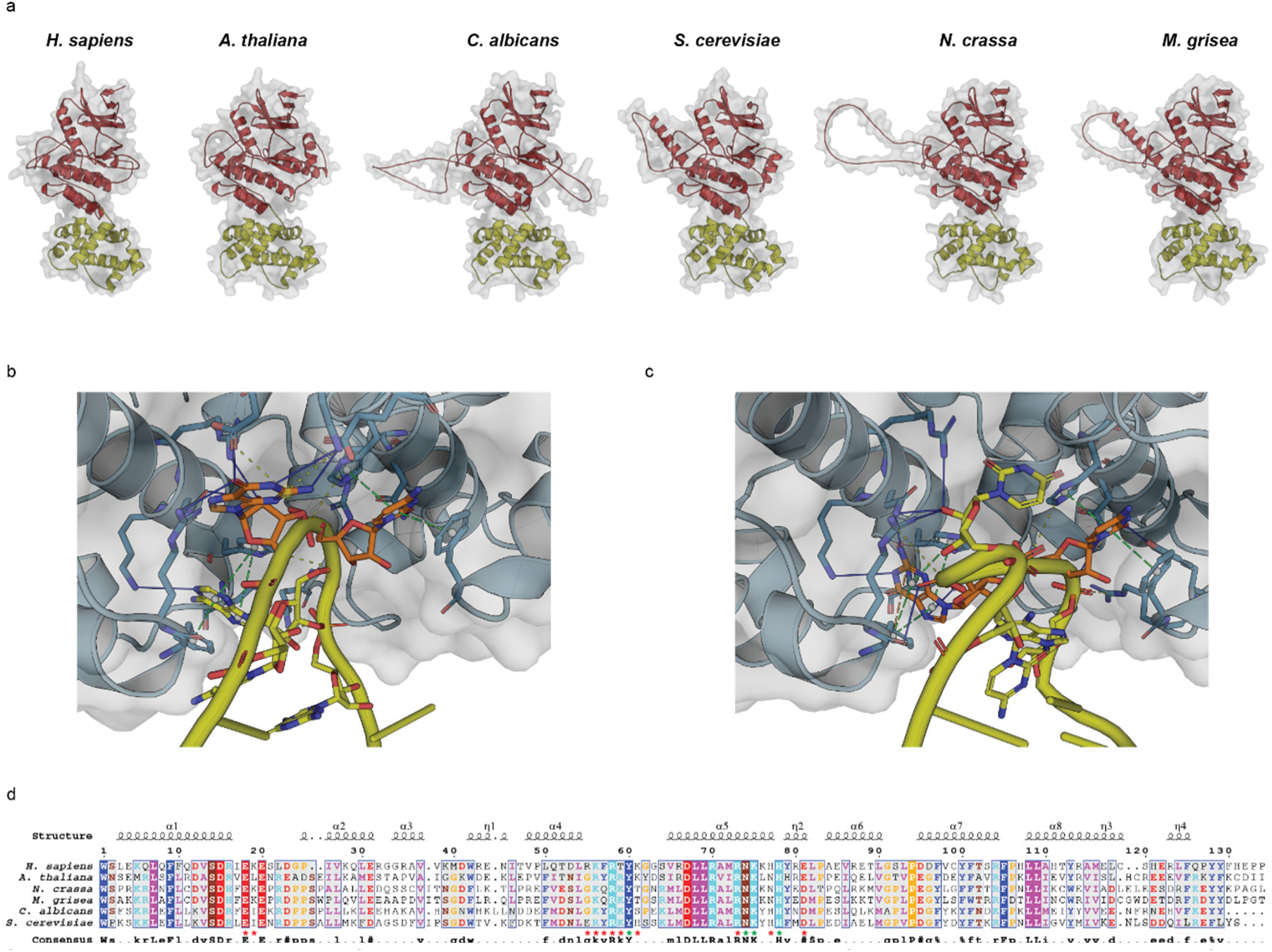
AlphaFold3 Predictions and Sequence Alignments of IRE1 Orthologues. **a,** AlphaFold3 predictions of the kinase (red) – RNase (yellow) regions for the IRE1 species used in the insulation screening (see Fig. 3), including *Homo sapiens*, *Arabidopsis thaliana*, *Candida albicans*, *Saccharomyces cerevisiae*, *Neurospora crassa*, and *Magnaporthe grisea*. **b,c,** AlphaFold3 predictions of the interaction between the 5’ loop (b), and the 3’ loop (c) of the XBP1 splicing region (in yellow) with the human RNase dimer of IRE1 (in light blue). The cleaved G and C bases are marked in orange. Amino acid residues of the RNase that interact with the loops are depicted as sticks, with the interactions highlighted by dashed lines. **d,** Sequence alignment of the RNase region from the chosen species. Asterisks identify positions predicted by AlphaFold3 in (b) and (c) to interact with the 5’ or 3’ XBP1 loop, with red denoting general interacting amino acids and green indicating previously identified catalytic amino acids. Consensus sequence (>65%) identity is shown below the alignment. Above the alignment, the secondary structure of the human IRE1 protein is depicted with α (alpha helix) and η (3_10_ helix) notation. Residues are colored according to their physicochemical properties: HKR (cyan), DE (red), STNQ (brown), AVLIM (pink), FYW (blue), PG (orange), and C (green).

**Supplementary Figure S3.**
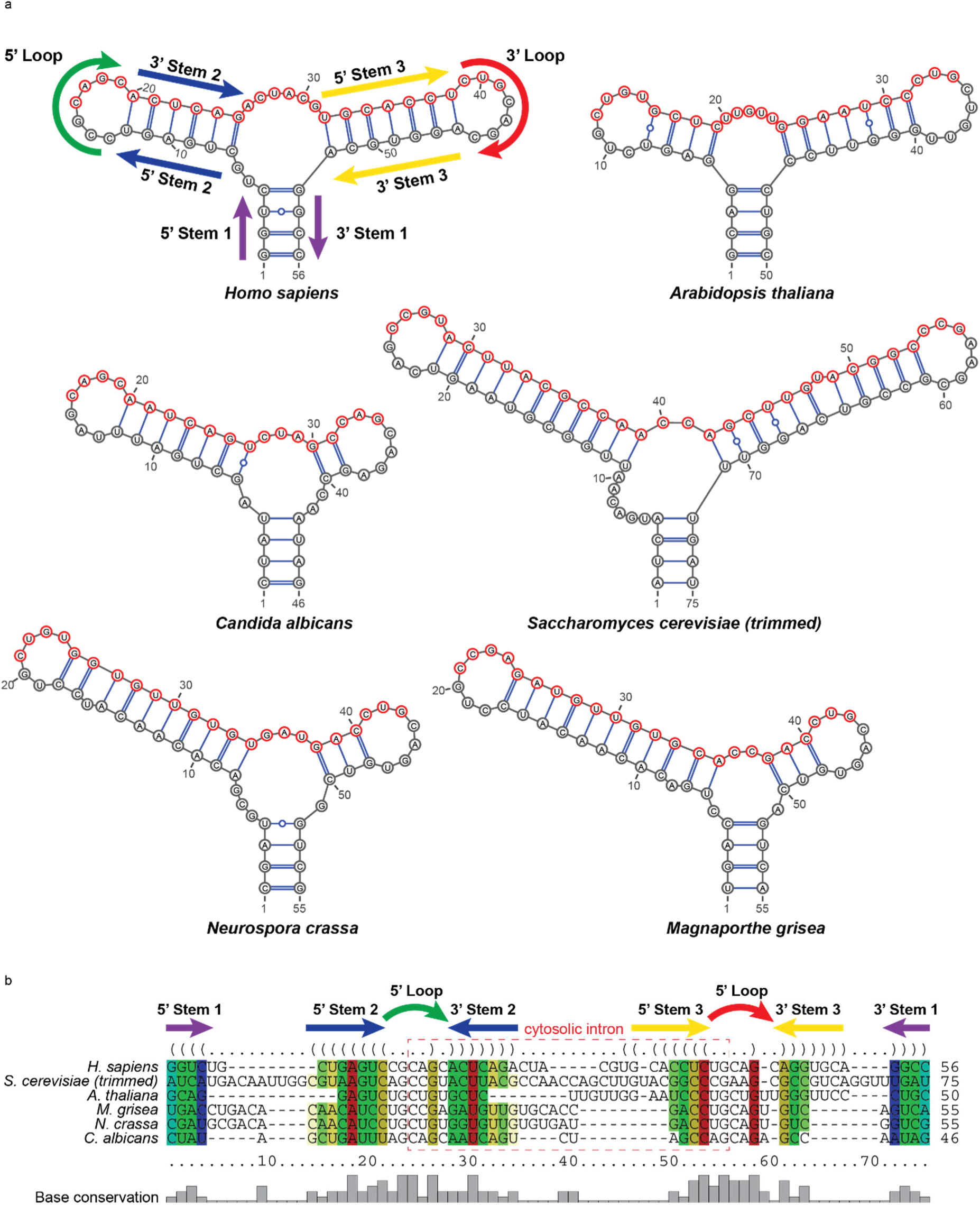
Structural Predictions and Sequence Alignments of the Bifurcated Stem-Loop Splicing Hairpin of different XBP1 Orthologues. **a,** mFold predictions of the secondary structure of the splicing hairpin region from *Homo sapiens*, *Arabidopsis thaliana*, *Candida albicans*, *Saccharomyces cerevisiae*, *Neurospora crassa*, and *Magnaporthe grisea* used for insulation screening (see Fig. 3). With the exception of the *Saccharomyces cerevisiae* sequence, in which the cytosolic intron was trimmed to reduce its length and to introduce a frameshift event after cytosolic splicing, all other sequences are endogenous. In the *Homo sapiens* XBP1 splicing hairpin, the conserved stem and loop structural features shared among all the orthologues are highlighted with different colored arrows. **b,** Sequence alignment of the XBP1 orthologues shown in (a). The sequence regions predicted to fold into conserved structural features are displayed on top, with arrows colored analogous to (a). For each position, the color scheme shows compatible base pairs where hue denotes the number of different types (1 = red, 2 = yellow, 3 = green, 4 = cyan, 5 = blue, and 6 = purple) of compatible base pairs (C-G, G-C, A-U, U-A, G-U, or U-G), and saturation decreases with the number of incompatible base pairs, thus showing sequence conservation within the column. Below the alignment, the histogram shows the frequency of the most conserved base pair for each position.

**Supplementary Figure S4.**
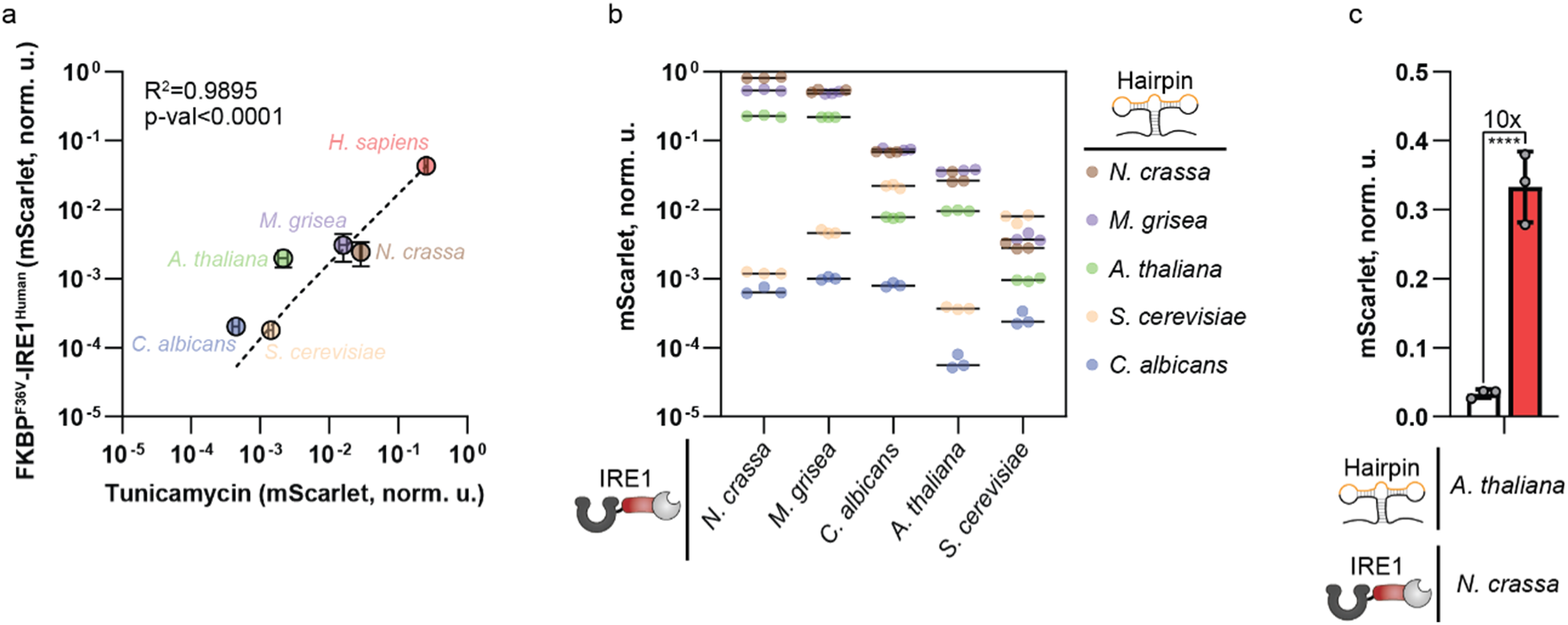
Functional Characterization of Orthologous IRE1 - Hairpin Pairs. **a,** Correlation between the splicing activity of endogenous IRE1 activated with 5 μg/mL of tunicamycin in HEK293 cells and the splicing activity of transfected FKBP^F36V^-IRE1^human^ activated with 500 nM of B/B homodimerizer in IRE1 HeLa KO cells used for the insulation screening. The plot shows mScarlet normalized units (norm. u.) obtained for each splicing reporter, with the species name color-coded to match the corresponding data point. The dashed line represents the curve fitting, with an R^2^ value of 0.9895 and a p < 0.0001 **b,** mScarlet normalized units (norm. u.) obtained from each splicing hairpin reporter species for each tested FKBP^F36V^-IRE1 ortholog. **c,** Dynamic range of the second-best insulating pair composed of the *Arabidopsis thaliana* hairpin and the *Neurospora crassa* IRE1. Individual data points are shown in gray, with error bars indicating the standard deviation from biological triplicates (n = 3). An unpaired t-test was used for statistical analysis, and asterisks indicate p < 0.0001 (****).

**Supplementary Figure S5.**
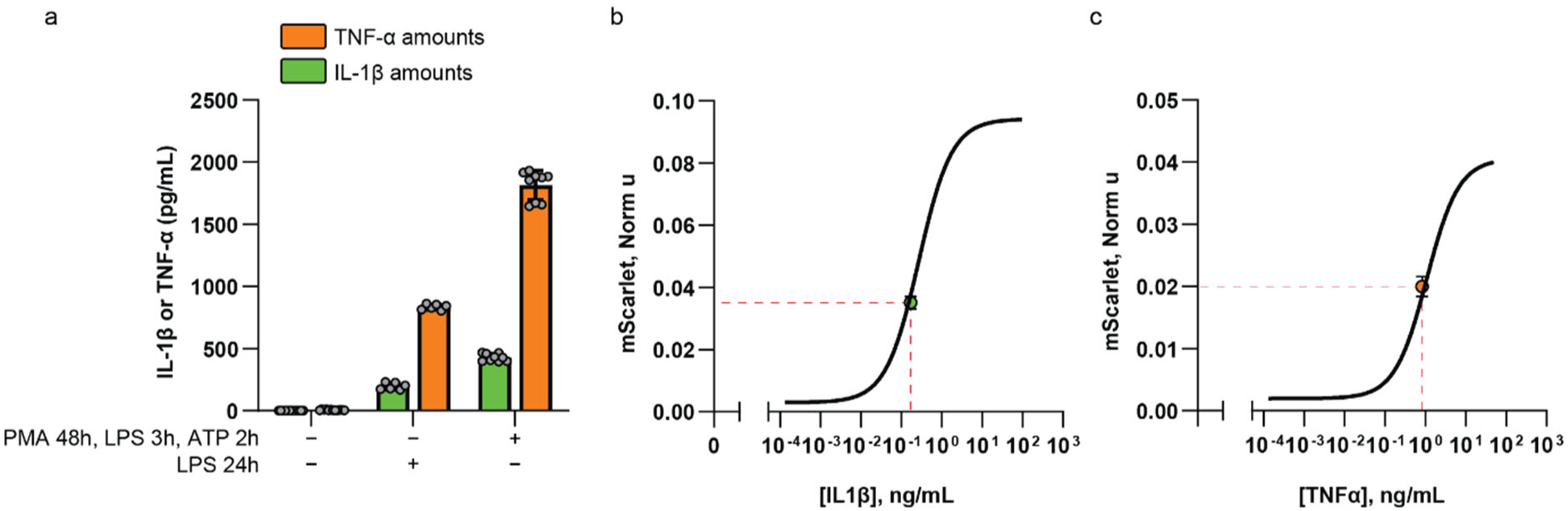
IL-1β and TNF-α Production by THP-1 Cells with Sensing by HEK293 Cells. **a,** Concentration of TNF-α and IL-1β (pg/mL) quantified by automated ELISA, produced by undifferentiated and unstimulated THP-1 cells, undifferentiated THP-1 cells treated overnight with 1 μg/mL of LPS, or macrophage-differentiated THP-1 cells stimulated with 1 μg/mL LPS for 3 hours (h) followed by 5 mM ATP for 2h. Shown in gray are individual data points, with error bars denoting the standard deviation from at least 6 biological replicates per condition (n ≥ 6). **b,** Average mScarlet signal (green dot, n = 3) of HEK293 cells transfected with IL-1R- and IL1RAcP-SMS mRNA upon addition of macrophage-differentiated and stimulated THP-1 medium on the SMS receptor. The signal is overlaid on the dose-response curve calculated with the purified IL-1β, and the concentration of IL-1β in the THP-1 medium was determined by automated ELISA. **c,** Average mScarlet signal (orange dot, n = 3) of HEK293 cells transfected with anti-TNF-α VHH-SMS mRNA co-cultured overnight with LPS-stimulated THP-1 monocytes. The signal is overlaid on the dose-response curve calculated with purified TNF-α, and the concentration of TNF-α in the co-culture medium was determined by automated ELISA.

**Supplementary Figure S6.**
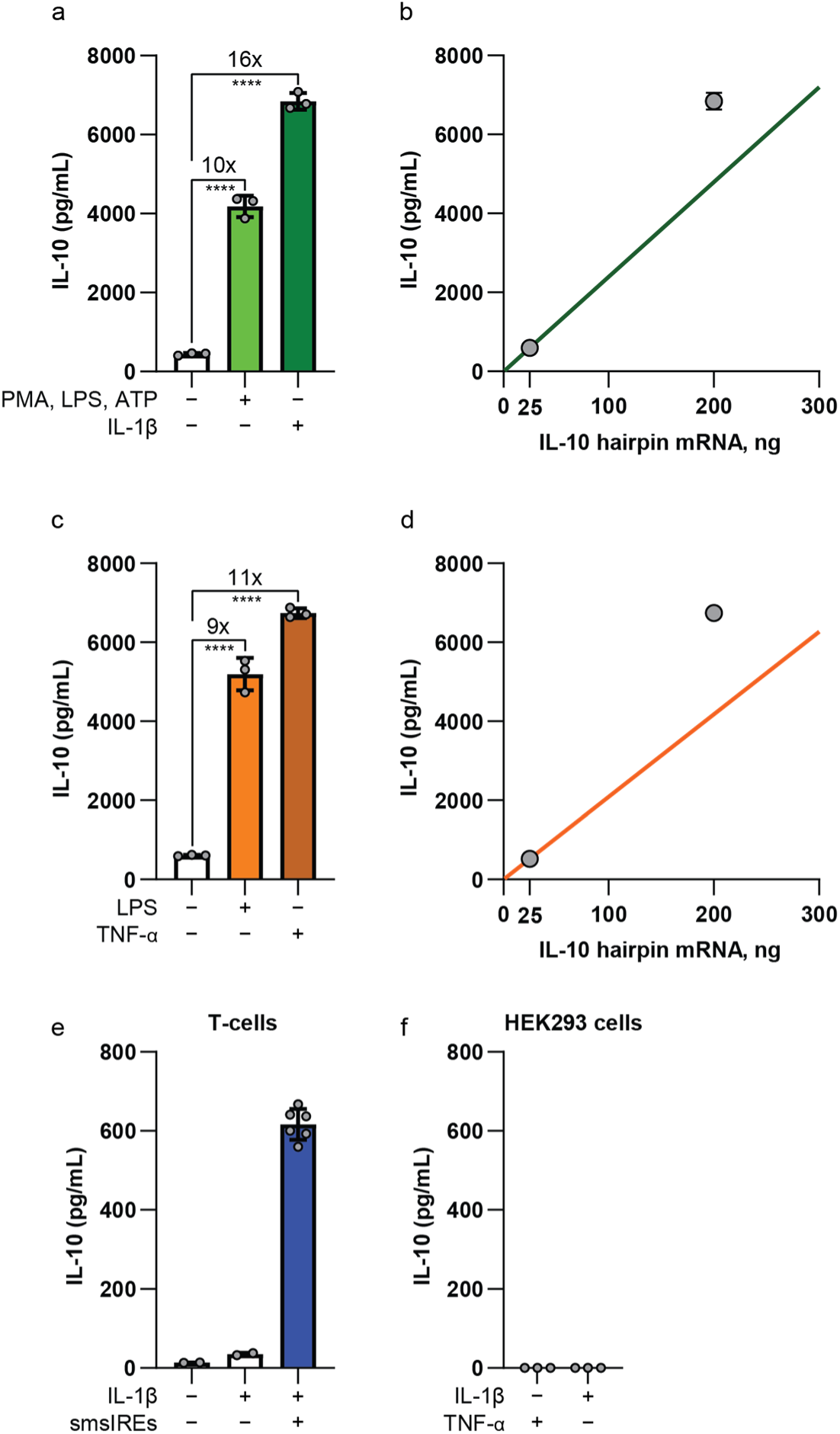
IL-10 Production with Varying Splicing Hairpin mRNA Amounts and Baseline IL-10 Expression in HEK293 Cells and Human Primary T-cells. **a,c,** IL-10 (pg/mL) released by HEK293 cells transfected with an 8x fold increase of the splicing hairpin mRNA for the IL-1β (a) or TNF-α (c) cytokine-converter SMS. For (a), IL-10 release was triggered either by the addition of macrophage-differentiated and stimulated THP-1 cell medium or by direct addition of purified IL-1β (10 ng/mL). For (c), IL-10 release was induced by overnight co-culture with LPS-stimulated THP-1 monocytes or by direct addition of purified TNF-α (10 ng/mL). Data are presented as the mean of biological triplicates (n = 3), with gray dot representing individual values and vertical bars indicating the standard deviation. ANOVA with Dunnett correction was performed to compare each condition to the control column, and asterisks indicate p < 0.0001 (****). **b,d,** Graphs showing IL-10 (pg/mL) production by HEK293 cells in response to sensing IL-1β (10 ng/mL, (b)) or TNF-α (10 ng/mL, (d)), with two different transfection amounts of IL-10 hairpin mRNA. The expected linear relationship between IL-10 production and mRNA quantity is indicated by the solid green or orange line, respectively. **e,** Endogenous IL-10 production levels by primary T-cells, either unstimulated or stimulated with IL-1β (n = 2), compared to IL-10 production by the active IL-1b cytokine-converter SMS receptor (n = 6). **f,** Endogenous IL-10 production by non-transfected HEK293 cells in the presence of IL-1β or TNF-α.

## Acknowledgements

This study was supported by ETH Zürich and by the Swiss National Science Foundation (SNSF), grant no. 182969. We are grateful to the Single Cell Unit of D-BSSE for their help with flow cytometry, cell sorting, and microscopy. We extend our thanks to Anthony Abraham, Bartolomeo Angelici, Vasileios Cheras, Jiten Doshi, Judith Johanna Jäckel, Philip Wolfgang Müller-Thümen, Gabriel Senn, David Schweingruber, and Fabian Trick for their insightful discussions.

## Author information

### Contributions

M.L. designed the study, performed experimental work, analyzed data and drafted the manuscript. Y.B. designed and supervised the study and revised the manuscript.

## Ethics declarations

### Competing interests

A patent application has been filed covering the technology described in this study. Y.B. is a shareholder and an employee of Pattern Biosciences.

