## Supplementary material for "Synthetic Cytosolic Splicing Enables Programmable mRNA-Encoded Receptors": See Table S1

| Plasmid ID & Features | Functional Sequence |
|---|---|
| pMLS001_(SplicingReporter$^{Hs}$)<br>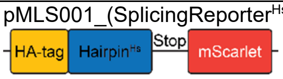 | GCCACCATGTATCCTTATGATGTTCCAGATTATGCTGGCCTTGTAGTTGAGAACCAGGAG<br>TTAAGACAGCGCTTGGGGATGGATGCCCTGGTTGCTGAAGAGGAGGCGGAAGCCAAGG<br>GGAATGAAGTGAGGCCAGTGGCCGGGTCTGCTGAGTCCGcagcactcagactacgtgcacctctgC<br>AGCAGGTGCAGGCCCAGTTGTCACCCCTCCAGAACATCTCCCCTTGGATTCTGGCGGTA<br>TTGACTCTTCAGATTCAGAGTCTGATATCCTGTTGGGCATTCTAATAGGATCCGTTTCAAA<br>AGGGGAAGCCGTCATTAAAGAATTTATGCGATTTAAGGTTCATATGGAAGGCTCTATGAA<br>TGGACATGAATTTGAAATTGAAGGGGAAGGGGAAGGGCGGCCATATGAAGGAACGCAAA<br>CGGCAAAATTGAAAGTCACAAAAGGCGGGCCGCTTCCCTTTAGCTGGGATATATTGTCAC<br>CACAATTTATGTATGGGAGCAGAGCTTTTACTAAACATCCTGCTGATATCCCAGATTATTA<br>CAAACAGTCATTTCCTGAAGGGTTTAAATGGGAACGGGTTATGAATTTTGAAGATGGTGG<br>GGCTGTAACTGTTACACAAGATACTAGCTTGGAAGACGGAACTCTTATATATAAAGTCAAA<br>CTTAGGGGAACAAATTTTCCACCCGATGGACCTGTTATGCAAAAGAAAACGATGGGTTGG<br>GAAGCAAGCACAGAACGACTTTATCCAGAAGATGGTGTTTTGAAAGGGGATATCAAAATG<br>GCTCTTCGGTTGAAAGATGGTGGTCGCTATCTCGCTGATTTCAAAACGACTTATAAAGCT<br>AAGAAACCAGTTCAAATGCCAGGGGCGTATAATGTAGATCGGAAACTGGATATTACATCT<br>CATAATGAAGATTATACTGTAGTTGAGCAATATGAGCGGAGCGAAGGGAGACATAGTACT<br>GGCGGAATGGATGAACTCTATAAATAG |
| pMLS002_(SplicingReporter$^{Hs}_{Spliced}$)<br>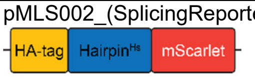 | GCCACCATGTATCCTTATGATGTTCCAGATTATGCTGGCCTTGTAGTTGAGAACCAGGAG<br>TTAAGACAGCGCTTGGGGATGGATGCCCTGGTTGCTGAAGAGGAGGCGGAAGCCAAGG<br>GGAATGAAGTGAGGCCAGTGGCCGGGTCTGCTGAGTCCGCAGCAGGTGCAGGCCCAGT<br>TGTCACCCCTCCAGAACATCTCCCCTTGGATTCTGGCGGTATTGACTCTTCAGATTCAGA<br>GTCTGATATCCTGTTGGGCATTCTAATAGGATCCGTTTCAAAAGGGGAAGCCGTCATTAA<br>AGAATTTATGCGATTTAAGGTTCATATGGAAGGCTCTATGAATGGACATGAATTTGAAATT<br>GAAGGGGAAGGGGAAGGGCGGCCATATGAAGGAACGCAAACGGCAAAATTGAAAGTCA<br>CAAAAGGCGGGCCGCTTCCCTTTAGCTGGGATATATTGTCACCACAATTTATGTATGGGA<br>GCAGAGCTTTTACTAAACATCCTGCTGATATCCCAGATTATTACAAACAGTCATTTCCTGA<br>AGGGTTTAAATGGGAACGGGTTATGAATTTTGAAGATGGTGGGGCTGTAACTGTTACACA<br>AGATACTAGCTTGGAAGACGGAACTCTTATATATAAAGTCAAACTTAGGGGAACAAATTTT<br>CCACCCGATGGACCTGTTATGCAAAAGAAAACGATGGGTTGGGAAGCAAGCACAGAACG<br>ACTTTATCCAGAAGATGGTGTTTTGAAAGGGGATATCAAAATGGCTCTTCGGTTGAAAGA<br>TGGTGGTCGCTATCTCGCTGATTTCAAAACGACTTATAAAGCTAAGAAACCAGTTCAAATG<br>CCAGGGGCGTATAATGTAGATCGGAAACTGGATATTACATCTCATAATGAAGATTATACT<br>GTAGTTGAGCAATATGAGCGGAGCGAAGGGAGACATAGTACTGGCGGAATGGATGAACT<br>CTATAAATAG |
| pMLS017_(SplicingReporter$^{Sc}_{Trimmed}$)<br>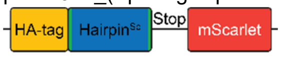 | GCCACCATGTATCCTTATGATGTTCCAGATTATGCTTACAATTCGAATGATCATGACAATT<br>GGCGTAAGTCAGccgtacttacgccaaccagcttgtacggcccgAAGCGCCGTCAGGTTTGATCATTCG<br>AATTGTAGGATCCGTTTCAAAAGGGGAAGCCGTCATTAAAGAATTTATGCGATTTAAGGTT<br>CATATGGAAGGCTCTATGAATGGACATGAATTTGAAATTGAAGGGGAAGGGGAAGGGCG<br>GCCATATGAAGGAACGCAAACGGCAAAATTGAAAGTCACAAAAGGCGGGCCGCTTCCCT<br>TTAGCTGGGATATATTGTCACCACAATTTATGTATGGGAGCAGAGCTTTTACTAAACATCC<br>TGCTGATATCCCAGATTATTACAAACAGTCATTTCCTGAAGGGTTTAAATGGGAACGGGTT<br>ATGAATTTTGAAGATGGTGGGGCTGTAACTGTTACACAAGATACTAGCTTGGAAGACGGA<br>ACTCTTATATATAAAGTCAAACTTAGGGGAACAAATTTTCCACCCGATGGACCTGTTATGC<br>AAAAGAAAACGATGGGTTGGGAAGCAAGCACAGAACGACTTTATCCAGAAGATGGTGTTT<br>TGAAAGGGGATATCAAAATGGCTCTTCGGTTGAAAGATGGTGGTCGCTATCTCGCTGATT<br>TCAAAACGACTTATAAAGCTAAGAAACCAGTTCAAATGCCAGGGGCGTATAATGTAGATC<br>GGAAACTGGATATTACATCTCATAATGAAGATTATACTGTAGTTGAGCAATATGAGCGGA<br>GCGAAGGGAGACATAGTACTGGCGGAATGGATGAACTCTATAAATAG |
| pMLS021_($_{Cytosolic}$IRE1$^{Hs}$)<br>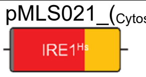 | MHQQQQLQHQQFQKELEKIQLLQQQQQQLPFHPPGDTAQDGELLDTSGPYSESSGTSSPS<br>TSPRASNHSLCSGSSASKAGSSPSLEQDDGDEETSVVIVGKISFCPKDVLGHGAEGTIVYRG<br>MFDNRDVAVKRILPECFSFADREVQLLRESDEHPNVIRYFCTEKDRQFQYIAIELCAATLQEYV<br>EQKDFAHLGLEPITLLQQTTSGLAHLHSLNIVHRDLKPHNILISMPNAHGKIKAMISDFGLCKKL<br>AVGRHSFSRRSGVPGTEGWIAPEMLSEDCKENPTYTVDIFSAGCVFYYVISEGSHPFGKSLQ<br>RQANILLGACSLDCLHPEKHEDVIARELIEKMIAMDPQKRPSAKHVLKHPFFWSLEKQLQFFQ<br>DVSDRIEKESLDGPIVKQLERGGRAVVKMDWRENITVPLQTDLRKFRTYKGGSVRDLLRAMR<br>NKKHHYRELPAEVRETLGSLPDDFVCYFTSRFPHLLAHTYRAMELCSHERLFQPYYFHEPPE<br>PQPPVTPDAL |
| pMLS022_(CCdimer-$_{Cytosolic}$IRE1$^{Hs}$)<br>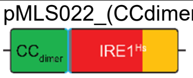 | MGEIAALKQEIAALKKENAALKWEIAALKQGYYGGSGGTGGSGGTSMHQQQQLQHQQFQK<br>ELEKIQLLQQQQQQLPFHPPGDTAQDGELLDTSGPYSESSGTSSPSTSPRASNHSLCSGSS<br>ASKAGSSPSLEQDDGDEETSVVIVGKISFCPKDVLGHGAEGTIVYRGMFDNRDVAVKRILPEC<br>FSFADREVQLLRESDEHPNVIRYFCTEKDRQFQYIAIELCAATLQEYVEQKDFAHLGLEPITLL<br>QQTTSGLAHLHSLNIVHRDLKPHNILISMPNAHGKIKAMISDFGLCKKLAVGRHSFSRRSGVP<br>GTEGWIAPEMLSEDCKENPTYTVDIFSAGCVFYYVISEGSHPFGKSLQRQANILLGACSLDCL<br>HPEKHEDVIARELIEKMIAMDPQKRPSAKHVLKHPFFWSLEKQLQFFQDVSDRIEKESLDGPI<br>VKQLERGGRAVVKMDWRENITVPLQTDLRKFRTYKGGSVRDLLRAMRNKKHHYRELPAEVR<br>ETLGSLPDDFVCYFTSRFPHLLAHTYRAMELCSHERLFQPYYFHEPPEPQPPVTPDAL |
| pMLS023_(HexCoilAla-$_{Cytosolic}$IRE1$^{Hs}$)<br>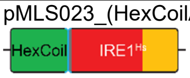 | MAEAESALEYAQQALEKAQLALQAARQALKAGGSGGTGGSGGTSMHQQQQLQHQQFQKE<br>LEKIQLLQQQQQQLPFHPPGDTAQDGELLDTSGPYSESSGTSSPSTSPRASNHSLCSGSSA<br>SKAGSSPSLEQDDGDEETSVVIVGKISFCPKDVLGHGAEGTIVYRGMFDNRDVAVKRILPECF<br>SFADREVQLLRESDEHPNVIRYFCTEKDRQFQYIAIELCAATLQEYVEQKDFAHLGLEPITLLQ<br>QTTSGLAHLHSLNIVHRDLKPHNILISMPNAHGKIKAMISDFGLCKKLAVGRHSFSRRSGVPGT<br>EGWIAPEMLSEDCKENPTYTVDIFSAGCVFYYVISEGSHPFGKSLQRQANILLGACSLDCLHP<br>EKHEDVIARELIEKMIAMDPQKRPSAKHVLKHPFFWSLEKQLQFFQDVSDRIEKESLDGPIVK<br>QLERGGRAVVKMDWRENITVPLQTDLRKFRTYKGGSVRDLLRAMRNKKHHYRELPAEVRET<br>LGSLPDDFVCYFTSRFPHLLAHTYRAMELCSHERLFQPYYFHEPPEPQPPVTPDAL |

| | |
|---|---|
| pMLS025_(I301-$_{\text{Cytosolic}}$IRE1$^{\text{Hs}}$)<br>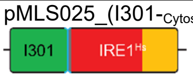 | MKMEELFKKHKIVAVLRANSVEEAKKKALAVFLGGVHLIEITFTVPDADTVIKELSFLKEMGAII<br>GAGTVTSVEQCRKAVESGAEFIVSPHLDEEISQFCKEKGVFYMPGVMTPTELVKAMKLGHTIL<br>KLFPGEVVGPQFVKAMKGPFPNVKFVPTGGVNLDNVCEWFKAGVLAVGVGSALVKGTPVEV<br>AEKAKAFVEKIRGCTEGGSGGTGGSGGTSMHQQQQLQHQQFQKELEKIQLLQQQQQQLPF<br>HPPGDTAQDGELLDTSGPYSESSGTSSPSTSPRASNHSLCSGSSASKAGSSPSLEQDDGDE<br>ETSVVIVGKISFCPKDVLGHGAEGTIVYRGMFDNRDVAVKRILPECFSFADREVQLLRESDEH<br>PNVIRYFCTEKDRQFQYIAIELCAATLQEYVEQKDFAHLGLEPITLLQQTTSGLAHLHSLNIVHR<br>DLKPHNILISMPNAHGKIKAMISDFGLCKKLAVGRHSFSRRSGVPGTEGWIAPEMLSEDCKEN<br>PTYTVDIFSAGCVFYYVISEGSHPFGKSLQRQANILLGACSLDCLHPEKHEDVIARELIEKMIAM<br>DPQKRPSAKHVLKHPFFWSLEKQLQFFQDVSDRIEKESLDGPIVKQLERGGRAVVKMDWRE<br>NITVPLQTDLRKFRTYKGGSVRDLLRAMRNKKHHYRELPAEVRETLGSLPDDFVCYFTSRFP<br>HLLAHTYRAMELCSHERLFQPYYFHEPPEPQPPVTPDAL |
| pMLS062_(FKBP-$_{\text{Cytosolic}}$IRE1$^{\text{Hs}}$)<br>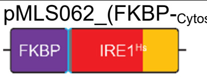 | MGVQVETISPGDGRTFPKRGQTCVVHYTGMLEDGKKFDSSRDRNKPFKFMLGKQEVIRGW<br>EEGVAQMSVGQRAKLTISPDYAYGATGHPGIIPPHATLVFDVELLKLEGGSGGTGGSGGTSM<br>HQQQQLQHQQFQKELEKIQLLQQQQQQLPFHPPGDTAQDGELLDTSGPYSESSGTSSPSTS<br>PRASNHSLCSGSSASKAGSSPSLEQDDGDEETSVVIVGKISFCPKDVLGHGAEGTIVYRGMF<br>DNRDVAVKRILPECFSFADREVQLLRESDEHPNVIRYFCTEKDRQFQYIAIELCAATLQEYVEQ<br>KDFAHLGLEPITLLQQTTSGLAHLHSLNIVHRDLKPHNILISMPNAHGKIKAMISDFGLCKKLAV<br>GRHSFSRRSGVPGTEGWIAPEMLSEDCKENPTYTVDIFSAGCVFYYVISEGSHPFGKSLQRQ<br>ANILLGACSLDCLHPEKHEDVIARELIEKMIAMDPQKRPSAKHVLKHPFFWSLEKQLQFFQDV<br>SDRIEKESLDGPIVKQLERGGRAVVKMDWRENITVPLQTDLRKFRTYKGGSVRDLLRAMRNK<br>KHHYRELPAEVRETLGSLPDDFVCYFTSRFPHLLAHTYRAMELCSHERLFQPYYFHEPPEPQ<br>PPVTPDAL |
| pMLS063_(FRB-$_{\text{Cytosolic}}$IRE1$^{\text{Hs}}$)<br>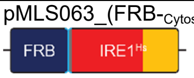 | MVAILWHEMWHEGLEEASRLYFGERNVKGMFEVLEPLHAMMERGPQTLKETSFNQAYGRD<br>LMEAQEWCRKYMKSGNVKDLLQAWDLYYHVFRRISGGSGGTGGSGGTSMHQQQQLQHQ<br>QFQKELEKIQLLQQQQQQLPFHPPGDTAQDGELLDTSGPYSESSGTSSPSTSPRASNHSLC<br>SGSSASKAGSSPSLEQDDGDEETSVVIVGKISFCPKDVLGHGAEGTIVYRGMFDNRDVAVKR<br>ILPECFSFADREVQLLRESDEHPNVIRYFCTEKDRQFQYIAIELCAATLQEYVEQKDFAHLGLE<br>PITLLQQTTSGLAHLHSLNIVHRDLKPHNILISMPNAHGKIKAMISDFGLCKKLAVGRHSFSRRS<br>GVPGTEGWIAPEMLSEDCKENPTYTVDIFSAGCVFYYVISEGSHPFGKSLQRQANILLGACSL<br>DCLHPEKHEDVIARELIEKMIAMDPQKRPSAKHVLKHPFFWSLEKQLQFFQDVSDRIEKESLD<br>GPIVKQLERGGRAVVKMDWRENITVPLQTDLRKFRTYKGGSVRDLLRAMRNKKHHYRELPA<br>EVRETLGSLPDDFVCYFTSRFPHLLAHTYRAMELCSHERLFQPYYFHEPPEPQPPVTPDAL |
| pMLS300_(FKBP$^{\text{F36V}}$-$_{\text{Cytosolic}}$IRE1$^{\text{Hs}}$)<br>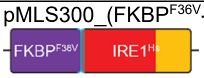 | MGVQVETISPGDGRTFPKRGQTCVVHYTGMLEDGKK**V**DSSRDRNKPFKFMLGKQEVIRGW<br>EEGVAQMSVGQRAKLTISPDYAYGATGHPGIIPPHATLVFDVELLKLEGGSGGTGGSGGTSM<br>HQQQQLQHQQFQKELEKIQLLQQQQQQLPFHPPGDTAQDGELLDTSGPYSESSGTSSPSTS<br>PRASNHSLCSGSSASKAGSSPSLEQDDGDEETSVVIVGKISFCPKDVLGHGAEGTIVYRGMF<br>DNRDVAVKRILPECFSFADREVQLLRESDEHPNVIRYFCTEKDRQFQYIAIELCAATLQEYVEQ<br>KDFAHLGLEPITLLQQTTSGLAHLHSLNIVHRDLKPHNILISMPNAHGKIKAMISDFGLCKKLAV<br>GRHSFSRRSGVPGTEGWIAPEMLSEDCKENPTYTVDIFSAGCVFYYVISEGSHPFGKSLQRQ<br>ANILLGACSLDCLHPEKHEDVIARELIEKMIAMDPQKRPSAKHVLKHPFFWSLEKQLQFFQDV<br>SDRIEKESLDGPIVKQLERGGRAVVKMDWRENITVPLQTDLRKFRTYKGGSVRDLLRAMRNK<br>KHHYRELPAEVRETLGSLPDDFVCYFTSRFPHLLAHTYRAMELCSHERLFQPYYFHEPPEPQ<br>PPVTPDAL |
| pMLS326_(SplicingReporter$^{\text{At}}$)<br>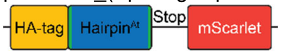 | GCCACCATGTATCCTTATGATGTTCCAGATTATGCTTACAATTCGAATGATAATACTACCA<br>TGATGTCGAAGCAGGAGTCTGctgtgctcttgttggaatccctgCTGTTGGGTTCCCTGCTTTGGCTT<br>CTGGGAGTAAAATCATTCGAATTGTAGGATCCGTTTCAAAAGGGGAAGCCGTCATTAAAG<br>AATTTATGCGATTTAAGGTTCATATGGAAGGCTCTATGAATGGACATGAATTTGAAATTGA<br>AGGGGAAGGGGAAGGGCGGCCATATGAAGGAACGCAAACGGCAAAATTGAAAGTCACA<br>AAAGGCGGGCCGCTTCCCTTTAGCTGGGATATATTGTCACCACAATTTATGTATGGGAGC<br>AGAGCTTTTACTAAACATCCTGCTGATATCCCAGATTATTACAAACAGTCATTTCCTGAAG<br>GGTTTAAATGGGAACGGGTTATGAATTTTGAAGATGGTGGGGCTGTAACTGTTACACAAG<br>ATACTAGCTTGGAAGACGGAACTCTTATATATAAAGTCAAACTTAGGGGAACAAATTTTCC<br>ACCCGATGGACCTGTTATGCAAAAGAAAACGATGGGTTGGGAAGCAAGCACAGAACGAC<br>TTTATCCAGAAGATGGTGTTTTGAAAGGGGATATCAAAATGGCTCTTCGGTTGAAAGATG<br>GTGGTCGCTATCTCGCTGATTTCAAAACGACTTATAAAGCTAAGAAACCAGTTCAAATGC<br>CAGGGGCGTATAATGTAGATCGGAAACTGGATATTACATCTCATAATGAAGATTATACTGT<br>AGTTGAGCAATATGAGCGGAGCGAAGGGAGACATAGTACTGGCGGAATGGATGAACTCT<br>ATAAATAG |
| pMLS327_(SplicingReporter$^{\text{Ca}}$)<br>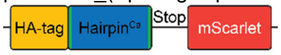 | GCCACCATGTATCCTTATGATGTTCCAGATTATGCTTACAATTCGAATGATTCCTCATCAT<br>CGTTAATTTCATCTTCCGGTCCAACTAATCATTCTATAGCTGATTTAGcagcaatcagtctagccag<br>CAGAGCCAATAGAATCATTTCAGTTGGACTTTGATGATTTTTTTCGATTTTAGGAATCATT<br>CGAATTGTAGGATCCGTTTCAAAAGGGGAAGCCGTCATTAAAGAATTTATGCGATTTAAG<br>GTTCATATGGAAGGCTCTATGAATGGACATGAATTTGAAATTGAAGGGGAAGGGGAAGG<br>GCGGCCATATGAAGGAACGCAAACGGCAAAATTGAAAGTCACAAAAGGCGGGCCGCTTC<br>CCTTTAGCTGGGATATATTGTCACCACAATTTATGTATGGGAGCAGAGCTTTTACTAAACA<br>TCCTGCTGATATCCCAGATTATTACAAACAGTCATTTCCTGAAGGGTTTAAATGGGAACG<br>GGTTATGAATTTTGAAGATGGTGGGGCTGTAACTGTTACACAAGATACTAGCTTGGAAGA<br>CGGAACTCTTATATATAAAGTCAAACTTAGGGGAACAAATTTTCCACCCGATGGACCTGTT<br>ATGCAAAAGAAAACGATGGGTTGGGAAGCAAGCACAGAACGACTTTATCCAGAAGATGG<br>TGTTTTGAAAGGGGATATCAAAATGGCTCTTCGGTTGAAAGATGGTGGTCGCTATCTCGC<br>TGATTTCAAAACGACTTATAAAGCTAAGAAACCAGTTCAAATGCCAGGGGCGTATAATGTA<br>GATCGGAAACTGGATATTACATCTCATAATGAAGATTATACTGTAGTTGAGCAATATGAGC<br>GGAGCGAAGGGAGACATAGTACTGGCGGAATGGATGAACTCTATAAATAG |
| pMLS329_(SplicingReporter$^{\text{Mg}}$)<br>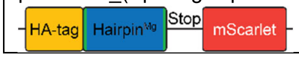 | GCCACCATGTATCCTTATGATGTTCCAGATTATGCTTACAATTCGAATGATGAAGCGAAGA<br>CCTCGGCTGACCTGACACAACATCCTGccgagatgttgtgcaccgacctgCAGTGTCAGTCAGCCA<br>AGGTCCCGCAAGCATCATTCGAATTGTAGGATCCGTTTCAAAAGGGGAAGCCGTCATTAA |

| | |
|---|---|
| | AGAATTTATGCGATTTAAGGTTCATATGGAAGGCTCTATGAATGGACATGAATTTGAAATT<br>GAAGGGGAAGGGGAAGGGCGGCCATATGAAGGAACGCAAACGGCAAAATTGAAAGTCA<br>CAAAAGGCGGGCCGCTTCCCTTTAGCTGGGATATATTGTCACCACAATTTATGTATGGGA<br>GCAGAGCTTTTACTAAACATCCTGCTGATATCCCAGATTATTACAAACAGTCATTTCCTGA<br>AGGGTTTAAATGGGAACGGGTTATGAATTTTGAAGATGGTGGGGCTGTAACTGTTACACA<br>AGATACTAGCTTGGAAGACGGAACTCTTATATATAAAGTCAAACTTAGGGGAACAAATTTT<br>CCACCCGATGGACCTGTTATGCAAAAGAAAACGATGGGTTGGGAAGCAAGCACAGAACG<br>ACTTTATCCAGAAGATGGTGTTTTGAAAGGGGATATCAAAATGGCTCTTCGGTTGAAAGA<br>TGGTGGTCGCTATCTCGCTGATTTCAAAACGACTTATAAAGCTAAGAAACCAGTTCAAATG<br>CCAGGGGCGTATAATGTAGATCGGAAACTGGATATTACATCTCATAATGAAGATTATACT<br>GTAGTTGAGCAATATGAGCGGAGCGAAGGGAGACATAGTACTGGCGGAATGGATGAACT<br>CTATAAATAG |
| pMLS330_(SplicingReporter$^{Nc}$) | GCCACCATGTATCCTTATGATGTTCCAGATTATGCTTACAATTCGAATGATGCGGATGCTA<br>CCTCCACTGCGACTGTGGACACTTCCCCCGATGCGACACAACATCCTGctgtggtgttgtgtgatg<br>acctgCAGTGTCGGTCGGCGGAAGTGCCACGCTCCAAGTGCCTGGCAGTGTCCCAATCAT<br>TCGAATTGTAGGATCCGTTTCAAAAGGGGAAGCCGTCATTAAAGAATTTATGCGATTTAA<br>GGTTCATATGGAAGGCTCTATGAATGGACATGAATTTGAAATTGAAGGGGAAGGGGAAG<br>GGCGGCCATATGAAGGAACGCAAACGGCAAAATTGAAAGTCACAAAAGGCGGGCCGCTT<br>CCCTTTAGCTGGGATATATTGTCACCACAATTTATGTATGGGAGCAGAGCTTTTACTAAAC<br>ATCCTGCTGATATCCCAGATTATTACAAACAGTCATTTCCTGAAGGGTTTAAATGGGAACG<br>GGTTATGAATTTTGAAGATGGTGGGGCTGTAACTGTTACACAAGATACTAGCTTGGAAGA<br>CGGAACTCTTATATATAAAGTCAAACTTAGGGGAACAAATTTTCCACCCGATGGACCTGTT<br>ATGCAAAAGAAAACGATGGGTTGGGAAGCAAGCACAGAACGACTTTATCCAGAAGATGG<br>TGTTTTGAAAGGGGATATCAAAATGGCTCTTCGGTTGAAAGATGGTGGTCGCTATCTCGC<br>TGATTTCAAAACGACTTATAAAGCTAAGAAACCAGTTCAAATGCCAGGGGCGTATAATGTA<br>GATCGGAAACTGGATATTACATCTCATAATGAAGATTATACTGTAGTTGAGCAATATGAGC<br>GGAGCGAAGGGAGACATAGTACTGGCGGAATGGATGAACTCTATAAATAG |
| pMLS380_(FKBP$^{F36V}$-$_{Cytosolic}$IRE1$^{Mg}$) | MGVQVETISPGDGRTFPKRGQTCVVHYTGMLEDGKK**V**DSSRDRNKPFKFMLGKQEVIRGW<br>EEGVAQMSVGQRAKLTISPDYAYGATGHPGIIPPHATLVFDVELLKLEGGSGGTGGSGGTSM<br>HGSVSRDDDPPQASVSEIVDKAKQLGDAPRRIEPDLRTIIDNVQDLTGPIYKMGSLEVNEDQQ<br>LGTGSNGTVVFAGKWDGRDVAVKRMLIQFYDIASQETRLLRESDDHPNVIRYYAQQSRDAFL<br>YIALELCQASLAEVIEKPAYFKNLAQAGEKDLPNVLYQITNGLSHLHSLRIVHRDLKPQNILVNM<br>GKDGKPRLLVSDFGLCKKLEGGQSSFGATTAHAAGTTGWRAPELLLDDDARDNTATMVDAS<br>MSSAHSGSGSVQGSSDVPNRRATRAIDIFSLGLVFFYVLTKGSHPYDRGDRYMREVNIRKGS<br>FDLSRLEVLGDYAMEARDIVERMLSFEPSERPTARDVMRHPFFWSAKKRLAFLCDVSDHFEK<br>EPRDPPSWPLQVLEEAAPDVITSGDFLRQLPREFVDSLGKQRKYTGSRMLDLLRALRNKKNH<br>YEDMPESLKKTVGPLPEGYLSFWTRRFDTLLINCWRIVIDCGWDETDRFRDYYDLPGT |
| pMLS381_(FKBP$^{F36V}$-$_{Cytosolic}$IRE1$^{Nc}$) | MGVQVETISPGDGRTFPKRGQTCVVHYTGMLEDGKK**V**DSSRDRNKPFKFMLGKQEVIRGW<br>EEGVAQMSVGQRAKLTISPDYAYGATGHPGIIPPHATLVFDVELLKLEGGSGGTGGSGGTSM<br>HTTDAPAEVVKPKRKAHRGRRGGIKHRKGPRNENTQSRDDEPPEPTVDEVVKKAQEIGQQP<br>KLEPDVITIPNGVDNVSGPILKMGSLEVNQEQQLGIGSNGTIVFAGKWDGRDVAVKRMLVQF<br>NEIASQETKLLRESDDHPNVIRYFAQQQSAGFLYIALELCQASLADVIQRPSMFRELAQAGER<br>DMPGVLYQVAKGLSHLHSLRIVHRDLKPQNILVNMGKDGRPRILVSDFGLCKKLEGGQSSFG<br>ATTAHAAGTTGWRAPELLLDDDGGPGPGATMTFTDPGSSMHSASGTGSGVVGAGVNVRRV<br>TRAIDIFSLGLVFFYVLTKGHHPFDLGDRYMRESNIRKGKYDLQLLEVLGDYAHDAKDLIESML<br>NSNPKKRPTAIGVMAHPFFWSPRKRLNFLCDVSDHFEKEPRDPPSPALALLEDQSSCVITNG<br>DFLKTLPREFVESLGKQRKYTGNRMLDLLRALRNKKNHYEDLTPQLRKMVGPLPEGYLGFFT<br>TRFPNLLIKCWEVIADLELEESDRFKEYYEPAGL |
| pMLS382_(FKBP$^{F36V}$-$_{TMD}$IRE1$^{Mg}$) | MCRAISLRRLLLLLLQLSQLLAVTQGMGVQVETISPGDGRTFPKRGQTCVVHYTGMLEDGKK<br>**V**DSSRDRNKPFKFMLGKQEVIRGWEEGVAQMSVGQRAKLTISPDYAYGATGHPGIIPPHATL<br>VFDVELLKLEGGSGGTGGSGGTGGSGGTGGSGGSSTITLIIFGVMAGVIGTILLISYGISMHGS<br>VSRDDDPPQASVSEIVDKAKQLGDAPRRIEPDLRTIIDNVQDLTGPIYKMGSLEVNEDQQLGT<br>GSNGTVVFAGKWDGRDVAVKRMLIQFYDIASQETRLLRESDDHPNVIRYYAQQSRDAFLYIA<br>LELCQASLAEVIEKPAYFKNLAQAGEKDLPNVLYQITNGLSHLHSLRIVHRDLKPQNILVNMGK<br>DGKPRLLVSDFGLCKKLEGGQSSFGATTAHAAGTTGWRAPELLLDDDARDNTATMVDASMS<br>SAHSGSGSVQGSSDVPNRRATRAIDIFSLGLVFFYVLTKGSHPYDRGDRYMREVNIRKGSFD<br>LSRLEVLGDYAMEARDIVERMLSFEPSERPTARDVMRHPFFWSAKKRLAFLCDVSDHFEKEP<br>RDPPSWPLQVLEEAAPDVITSGDFLRQLPREFVDSLGKQRKYTGSRMLDLLRALRNKKNHYE<br>DMPESLKKTVGPLPEGYLSFWTRRFDTLLINCWRIVIDCGWDETDRFRDYYDLPGT |
| pMLS394_(IL1R$^{Hs}$-$_{TMD}$IRE1$^{Mg}$) | MKVLLRLICFIALLISSLEADKCKEREEKIILVSSANEIDVRPCPLNPNEHKGTITWYKDDSKTPV<br>STEQASRIHQHKEKLWFVPAKVEDSGHYYCVVRNSSYCLRIKISAKFVENEPNLCYNAQAIFK<br>QKLPVAGDGGLVCPYMEFFKNENNELPKLQWYKDCKPLLLDNIHFSGVKDRLIVMNVAEKHR<br>GNYTCHASYTYLGKQYPITRVIEFITLEENKPTRPVIVSPANETMEVDLGSQIQLICNVTGQLSD<br>IAYWKWNGSVIDEDDPVLGEDYYSVENPANKRRSTLITVLNISEIESRFYKHPFTCFAKNTHGI<br>DAAYIQLIYPVTNFQKSTITLIIFGVMAGVIGTILLISYGISMHGSVSRDDDPPQASVSEIVDKAK<br>QLGDAPRRIEPDLRTIIDNVQDLTGPIYKMGSLEVNEDQQLGTGSNGTVVFAGKWDGRDVAV<br>KRMLIQFYDIASQETRLLRESDDHPNVIRYYAQQSRDAFLYIALELCQASLAEVIEKPAYFKNLA<br>QAGEKDLPNVLYQITNGLSHLHSLRIVHRDLKPQNILVNMGKDGKPRLLVSDFGLCKKLEGGQ<br>SSFGATTAHAAGTTGWRAPELLLDDDARDNTATMVDASMSSAHSGSGSVQGSSDVPNRRA<br>TRAIDIFSLGLVFFYVLTKGSHPYDRGDRYMREVNIRKGSFDLSRLEVLGDYAMEARDIVERM<br>LSFEPSERPTARDVMRHPFFWSAKKRLAFLCDVSDHFEKEPRDPPSWPLQVLEEAAPDVITS<br>GDFLRQLPREFVDSLGKQRKYTGSRMLDLLRALRNKKNHYEDMPESLKKTVGPLPEGYLSF<br>WTRRFDTLLINCWRIVIDCGWDETDRFRDYYDLPGT |
| pMLS395_(IL1RAcP$^{Hs}$-$_{TMD}$IRE1$^{Mg}$) | MTLLWCVVSLYFYGILQSDASERCDDWGLDTMRQIQVFEDEPARIKCPLFEHFLKFNYSTAH<br>SAGLTLIWYWTRQDRDLEEPINFRLPENRISKEKDVLWFRPTLLNDTGNYTCMLRNTTYCSKV<br>AFPLEVVQKDSCFNSPMKLPVHKLYIEYGIQRITCPNVDGYFPSSVKPTITWYMGCYKIQNFN<br>NVIPEGMNLSFLIALISNNGNYTCVVTYPENGRTFHLTRTLTVKVVGSPKNAVPPVIHSPNDHV |

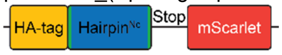


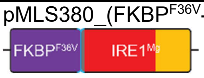


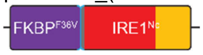


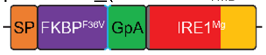


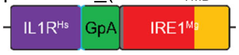


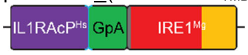

| | |
|---|---|
| | VYEKEPGEELLIPCTVYFSFLMDSRNEVWWTIDGKKPDDITIDVTINESISHSRTEDETRTQILSI<br>KKVTSEDLKRSYVCHARSAKGEVAKAAKVKQKVPAPRYTVESTITLIIFGVMAGVIGTILLISYGI<br>SMHGSVSRDDDPPQASVSEIVDKAKQLGDAPRRIEPDLRTIIDNVQDLTGPIYKMGSLEVNED<br>QQLGTGSNGTVVFAGKWDGRDVAVKRMLIQFYDIASQETRLLRESDDHPNVIRYYAQQSRD<br>AFLYIALELCQASLAEVIEKPAYFKNLAQAGEKDLPNVLYQITNGLSHLHSLRIVHRDLKPQNIL<br>VNMGKDGKPRLLVSDFGLCKKLEGGQSSFGATTAHAAGTTGWRAPELLLDDDARDNTATM<br>VDASMSSAHSGSGSVQGSSDVPNRRATRAIDIFSLGLVFFYVLTKGSHPYDRGDRYMREVNI<br>RKGSFDLSRLEVLGDYAMEARDIVERMLSFEPSERPTARDVMRHPFFWSAKKRLAFLCDVS<br>DHFEKEPRDPPSWPLQVLEEAAPDVITSGDFLRQLPREFVDSLGKQRKYTGSRMLDLLRALR<br>NKKNHYEDMPESLKKTVGPLPEGYLSFWTRRFDTLLINCWRIVIDCGWDETDRFRDYYDLPG<br>T |
| pMLS420_($\mathrm{FKBP}^{F36V}\text{-}{}_{Cytosolic}\mathrm{IRE1}^{At}$)<br>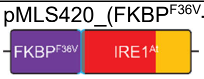 | MGVQVETISPGDGRTFPKRGQTCVVHYTGMLEDGKK**V**DSSRDRNKPFKFMLGKQEVIRGW<br>EEGVAQMSVGQRAKLTISPDYAYGATGHPGIIPPHATLVFDVELLKLEGGSGGTGGSGGTSM<br>HSSRGSDVSLKAGPSKKKKNRKSAKDTNRQSVPRGQDQFELIEGGQMLLGFNNFQSGATD<br>GRKIGKLFLSSKEIAKGSNGTVVFEGIYEGRPVAVKRLVRSHHEVAFKEIQNLIASDQHTNIIRW<br>YGVEYDQDFVYLSLERCTCSLDDLIKSYLEFSMTKVLENNDSTEGVAAYKIQLDSLEGVIKGN<br>NFWKVGGHPSPLMLKLMRDIVCGIVHLHELGIVHRDLKPQNVLISKDMTLSAKLSDMGISKRM<br>SRDMSSLGHLATGSGSSGWQAPEQLLQGRQTRAVDMFSLGCVIFYTITGCKHPFGDDLERD<br>VNIVKNKVDLFLVEHVPEASDLISRLLNPDPDLRPSATEVLLHPMFWNSEMRLSFLRDASDRV<br>ELENREADSEILKAMESTAPVAIGGKWDEKLEPVFITNIGRYRRYKYDSIRDLLRVIRNKLNHH<br>RELPPEIQELVGTVPEGFDEYFAVRFPKLLIEVYRVISLHCREEEVFRKYFKCDII |
| pMLS421_($\mathrm{FKBP}^{F36V}\text{-}{}_{Cytosolic}\mathrm{IRE1}^{Ca}$)<br>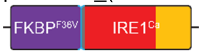 | MGVQVETISPGDGRTFPKRGQTCVVHYTGMLEDGKK**V**DSSRDRNKPFKFMLGKQEVIRGW<br>EEGVAQMSVGQRAKLTISPDYAYGATGHPGIIPPHATLVFDVELLKLEGGSGGTGGSGGTSM<br>HTTPKKKRKRGSRGGRRGGARKNNKQKTNGDEDQEDQQQNDESVVDEEIIPTKSLIPPSSL<br>PAIKSRKKLQIENNLVISDKILGYGSHGTVVFQGTFENRPVAVKRMLLDFYDIANHEVRLLQES<br>DDHPNVVRYFCSQSSESEKFLYIALELCLCTLEDIIEKPQNMPNLCIPKRNDILYQLTSGLHYLH<br>SLKIVHRDIKPQNILVANIKKNGKRKNQITEIDETCENNVRLLISDFGLCKKLENDQSSFRATTQ<br>NAASGTSGWRAPELLLNHDLWEISADSISSIHSNSNSNGNGNGNGATNGSVSNSATSGKRLT<br>KAIDIFSLGCVFYYILTGGYHPFGDRYLREGNIIKGEYDLSLLMEKCPNDRYESIDLISKIISHDP<br>SQRPNTGKILKHPLFWSFSKRLEFLLKVSDRFEIEKRDPPSPLLLKLEEHAKAVHNGNWHKLL<br>NDDEFMDNLGKYRKYSPEKLMDLLRAMRNKYHHYNDMPESLQLKMAPLPDGFYKYFNDKF<br>PKLLMEIYYVVEENFRNEHVFKEYY |
| pMLS422_($\mathrm{FKBP}^{F36V}\text{-}{}_{Cytosolic}\mathrm{IRE1}^{Sc}$)<br>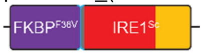 | MGVQVETISPGDGRTFPKRGQTCVVHYTGMLEDGKK**V**DSSRDRNKPFKFMLGKQEVIRGW<br>EEGVAQMSVGQRAKLTISPDYAYGATGHPGIIPPHATLVFDVELLKLEGGSGGTGGSGGTSM<br>HQRFKILPPLYVLLSKIGFMPEKEIPIVESKSLNCPSSSENVTKPFDMKSGKQVVFEGAVNDGS<br>LKSEKDNDDADEDDEKSLDLTTEKKKRKRGSRGGKKGRKSRIANIPNFEQSLKNLVVSEKILG<br>YGSSGTVVFQGSFQGRPVAVKRMLIDFCDIALMEIKLLTESDDHPNVIRYYCSETTDRFLYIAL<br>ELCNLNLQDLVESKNVSDENLKLQKEYNPISLLRQIASGVAHLHSLKIIHRDLKPQNILVSTSSR<br>FTADQQTGAENLRILISDFGLCKKLDSGQSSFRTNLNNPSGTSGWRAPELLEESNNLQCQVE<br>TEHSSSRHTVVSSDSFYDPFTKRRLTRSIDIFSMGCVFYYILSKGKHPFGDKYSRESNIIRGIFS<br>LDEMKCLHDRSLIAEATDLISQMIDHDPLKRPTAMKVLRHPLFWPKSKKLEFLLKVSDRLEIEN<br>RDPPSALLMKFDAGSDFVIPSGDWTVKFDKTFMDNLERYRKYHSSKLMDLLRALRNKYHHF<br>MDLPEDIAELMGPVPDGFYDYFTKRFPNLLIGVYMIVKENLSDDQILREFLYS |
| pMLS433_($\mathrm{LaG16\text{-}LaG2\text{-}}{}_{TMD}\mathrm{IRE1}^{Mg}$)<br>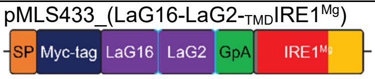 | MALPVTALLLPLALLLHAARPEQKLISEEDLMAQVQLVESGGRLVQAGDSLRLSCAASGRTFS<br>TSAMAWFRQAPGREREFVAAITWTVGNTILGDSVKGRFTISRDRAKNTVDLQMDNLEPEDTA<br>VYYCSARSRGYVLSVLRSVDSYDYWGQGTQVTVSGGGGSGGGGSGGGGSMAQVQLVES<br>GGGLVQAGGSLRLSCAASGRTFSNYAMGWFRQAPGKEREFVAAISWTGVSTYYADSVKGR<br>FTISRDNDKNTVYVQMNSLIPEDTAIYYCAAVRARSFSDTYSRVNEYDYWGQGTQVTVGGG<br>GSGGGGSGGGGSTGGSGGTGGSGGTGGSGGSSTGGSGGTGGSGGTSTITLIIFGVMAGVI<br>GTILLISYGISMHGSVSRDDDPPQASVSEIVDKAKQLGDAPRRIEPDLRTIIDNVQDLTGPIYKM<br>GSLEVNEDQQLGTGSNGTVVFAGKWDGRDVAVKRMLIQFYDIASQETRLLRESDDHPNVIRY<br>YAQQSRDAFLYIALELCQASLAEVIEKPAYFKNLAQAGEKDLPNVLYQITNGLSHLHSLRIVHR<br>DLKPQNILVNMGKDGKPRLLVSDFGLCKKLEGGQSSFGATTAHAAGTTGWRAPELLLDDDA<br>RDNTATMVDASMSSAHSGSGSVQGSSDVPNRRATRAIDIFSLGLVFFYVLTKGSHPYDRGD<br>RYMREVNIRKGSFDLSRLEVLGDYAMEARDIVERMLSFEPSERPTARDVMRHPFFWSAKKR<br>LAFLCDVSDHFEKEPRDPPSWPLQVLEEAAPDVITSGDFLRQLPREFVDSLGKQRKYTGSRM<br>LDLLRALRNKKNHYEDMPESLKKTVGPLPEGYLSFWTRRFDTLLINCWRIVIDCGWDETDRF<br>RDYYDLPGT |
| pMLS470_($\mathrm{SplicingHairpin}^{At}\text{-}{}_{T2A}\mathrm{IL10}^{Hs}$)<br>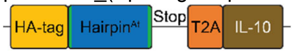 | GCCACCATGTATCCTTATGATGTTCCAGATTATGCTTACAATTCGAATGATAATACTACCA<br>TGATGTCGAAGCAGGAGTCTGctgtgctcttgttggaatccctgCTGTTGGGTTCCCTGCTTTGGCTT<br>CTGGGAGTAAAATCATTCGAATTGTAGGATCCGGAGAGGGCAGAGGAAGTCTGCTAACA<br>TGCGGTGACGTCGAGGAGAATCCTGGCCCACATTCTTCTGCACTCTTGTGTTGTCTCGTC<br>CTTTTAACTGGGGTCCGAGCCAGTCCGGGGCAAGGTACTCAATCAGAGAATAGTTGCAC<br>CCATTTTCCCGGCAACCTGCCGAATATGTTAAGGGACTTGAGGGATGCTTTCTCTCGGGT<br>CAAGACCTTCTTCCAAATGAAGGACCAACTCGACAACTTGCTGCTCAAAGAATCCCTGTT<br>AGAAGATTTTAAGGGGTATCTTGGTTGTCAAGCACTGTCAGAGATGATCCAGTTTTATTTA<br>GAGGAAGTTATGCCGCAAGCAGAAAACCAAGACCCAGATATAAAGGCGCACGTCAACTC<br>CCTTGGAGAAAACCTGAAAACTCTTCGACTGCGCCTGCGCAGGTGCCATAGATTTCTGC<br>CGTGCGAGAACAAAAGCAAAGCTGTGGAGCAGGTCAAGAATGCCTTCAATAAGCTGCAG<br>GAGAAGGGTATCTACAAAGCGATGTCTGAATTTGATATCTTCATAAACTATATCGAAGCCT<br>ATATGACTATGAAAATACGGAATTAG |
| pMLS479_($\mathrm{VHH2\text{-}}{}_{TMD}\mathrm{IRE1}^{Mg}$)<br>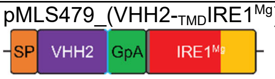 | MALPVTALLLPLALLLHAARPEQKLISEEDLMAQVQLVESGGGLVQPGGSLRLSCAASGFTFS<br>NYWMYWVRQAPGKGLEWVSEINTNGLITKYPDSVKGRFTISRDNAKNTLYLQMNSLKPEDT<br>ALYYCARSPSGFNRGQGTQVTVSSGGGGSGGGGSGGGGSTGGSGGTGGSGGTGGSGGT<br>GGSGGTGGSGGTSTITLIIFGVMAGVIGTILLISYGISMHGSVSRDDDPPQASVSEIVDKAKQL<br>GDAPRRIEPDLRTIIDNVQDLTGPIYKMGSLEVNEDQQLGTGSNGTVVFAGKWDGRDVAVKR<br>MLIQFYDIASQETRLLRESDDHPNVIRYYAQQSRDAFLYIALELCQASLAEVIEKPAYFKNLAQ |

| | |
|---|---|
| | AGEKDLPNVLYQITNGLSHLHSLRIVHRDLKPQNILVNMGKDGKPRLLVSDFGLCKKLEGGQS<br>SFGATTAHAAGTTGWRAPELLLDDDARDNTATMVDASMSSAHSGSGSVQGSSDVPNRRAT<br>RAIDIFSLGLVFFYVLTKGSHPYDRGDRYMREVNIRKGSFDLSRLEVLGDYAMEARDIVERML<br>SFEPSERPTARDVMRHPFFWSAKKRLAFLCDVSDHFEKEPRDPPSWPLQVLEEAAPDVITS<br>GDFLRQLPREFVDSLGKQRKYTGSRMLDLLRALRNKKNHYEDMPESLKKTVGPLPEGYLSF<br>WTRRFDTLLINCWRIVIDCGWDETDRFRDYYDLPGT |
